# Machine learning reveals signatures of promiscuous microbial amidases for micropollutant biotransformations

**DOI:** 10.1101/2024.08.09.606993

**Authors:** Thierry D. Marti, Diana Schweizer, Yaochun Yu, Milo R. Schärer, Silke I. Probst, Serina L. Robinson

## Abstract

Organic micropollutants - including pharmaceuticals, personal care products, pesticides and food additives - are prevalent in the environment and have unknown and potentially toxic effects. Humans are a direct source of micropollutants as the majority of pharmaceuticals are primarily excreted through urine. Urine contains its own microbiota with the potential to catalyze micropollutant biotransformations. Amidase signature (AS) enzymes are known for their promiscuous activity in micropollutant biotransformations, but the potential for AS enzymes from the urinary microbiota to transform micropollutants is not known. Moreover, characterization of AS enzymes to identify key chemical and enzymatic features predictive of biotransformation profiles is critical for developing benign-by-design chemicals and micropollutant removal strategies.

In this study, we biochemically characterized a new AS enzyme with arylamidase activity from a urine isolate, *Lacticaseibacillus rhamnosus,* and demonstrated its capability to hydrolyze pharmaceuticals and other micropollutants. To uncover the signatures of AS enzyme-substrate specificity, we then designed a targeted enzyme library consisting of 40 arylamidase homologs from diverse urine isolates and tested it against 17 structurally diverse compounds. We found that 16 out of the 40 enzymes showed activity on at least one substrate and exhibited diverse substrate specificities, with the most promiscuous enzymes active on nine different substrates.

Using an interpretable gradient boosting machine learning model, we identified chemical and amino acid features predictive of arylamidase biotransformations. Key chemical features from our substrates included the molecular weight of the amide carbonyl substituent and the number of charges in the molecule. Important amino acid features were found to be located on the protein surface and four predictive residues were located in close proximity of the substrate tunnel entrance.

Overall, this work highlights the understudied role of urine-derived microbial arylamidases and contributes to enzyme sequence-structure-substrate-based predictions of micropollutant biotransformations.

## Introduction

Micropollutants are environmental contaminants occurring at low concentrations, including pesticides, personal care products, industrial chemicals and pharmaceuticals. Two thirds of pharmaceuticals are excreted in urine, thus it is a major contributor of micropollutants in wastewater streams.^1^ Urine supplies the majority of nitrogen and phosphorus to wastewater streams^2–4^ and therefore has potential uses for resource recovery.^5,6^ However, many micropollutants have limited biodegradability and are therefore not effectively removed from urine through common procedures such as anaerobic storage^7–9^ or standard biological treatments,^10^ thereby hampering the applications of urine due to micropollutant contamination. Contrary to previous belief, urine itself is not sterile, and has its own native microbiota.^11–13^ Microbial biotransformations are considered as one of the dominant pathways for organic micropollutant removal,^14^ however, we lack knowledge of the capabilities of the urinary microbiota to biotransform micropollutants.

The nutrient-limited environment of the urinary tract, along with the significant exposure to micropollutants excreted in urine, may select for bacteria with the ability to degrade these micropollutants to use them as a carbon or energy source.^15^ Previously, we showed that urinary bacterial genomes encode homologs of enzymes involved in micropollutant biotransformations.^15^ Among 20 different enzyme classes investigated, the most prevalent enzyme class encoded in urinary bacterial genomes belonged to the amidase signature (AS) family. AS enzymes are of high relevance for micropollutant removal since they hydrolyze compounds such as propanil, chlorpropham, propham and paracetamol,^16^ the latter being one of the most abundant pharmaceuticals found in urine.^17^ AS enzymes are veritable ‘Swiss army knives’ capable of hydrolyzing multiple micropollutants including aliphatic, aromatic and branched amides and esters.^18^ However, fine-grained prediction capabilities for micropollutant biotransformations by AS enzymes are limited, even with the use of state-of-the-art enzyme-substrate prediction tools.^19,20^

AS enzymes are defined by a conserved stretch of ∼160 amino acids, termed the ‘amidase signature region’ that includes the catalytic triad composed of two serine and one lysine residue, and a glycine-serine-rich motif.^21^ A distinct subset of amidase signature enzymes investigated here are the aryl acylamidases (assigned to the Enzyme Commission number 3.5.1.13) which hydrolyze non-peptidic amide bonds typically producing aniline and a carboxylic acid anion.^22,23^ Under specific reaction conditions, aryl acylamidases can also catalyze the reverse reaction for amide bond formation,^22^ e.g., for the biomanufacturing of paracetamol (acetaminophen). Microbial biocatalysis has been employed for paracetamol production at the gram scale.^24^

Despite their medical and environmental relevance with potential biotechnological applications, acyl arylamidases from the urinary microbiota have not been biochemically characterized. Additionally, this study experimentally validated AS enzymes mined from genomic data to address the current challenge of predicting AS enzyme activity and promiscuity from sequencing data alone. Specifically, we performed chemical analysis of biotransformations catalyzed by a library of 40 urine-derived aryl acylamidases. We trained interpretable machine learning models to extract physicochemical enzyme and substrate features to uncover new micropollutant biotransformation relationships. More broadly, this work aims to inform enzyme engineering strategies, green chemical design, and biotransformation prediction.

## Methods

### Construction of amidase expression vectors for initial screening

Previous analyses^15^ revealed urinary bacterial genomes encode homologs to the paracetamol amidase from *Ochrobactrum* PP-2 *mah* (ANS81375.1)^15,16^ To experimentally test these homologs for paracetamol amidase activity, we selected three homologs here termed P203 = NCBI accession PKZ25206.1, P204 = NCBI accession PKZ66103.1, and P205 = NCBI accession PLA55254.1. The gene sequences were codon-optimized for expression in *Escherichia coli* using the Integrated DNA Technologies (IDT) codon optimization tool and synthesized as gBlocks (IDT). A C-terminal tobacco etch virus (TEV) site and 6XHis-tag along with homology arms to the MCS1 site of pCDFDuet-1 for Gibson assembly were added to the sequence. The amidases were cloned into MCS1 of pCDFDuet-1 using Gibson assembly kit (New England Biolabs) after linearization of the vector with NcoI and HindIII (Fermentas). The assembled plasmids were transformed in BL21(DE3) and T7 Express *E. coli* cells (New England Biolabs) by chemical transformation. Subsequently, 37 additional homologs of P205 were codon-optimized for *E. coli* using the build optimization software tool, BOOST,^25^ and cloned by the U.S. Department of Energy Joint Genome Institute with C-terminal 6x-His tags, a TEV cleavage site, and additional flexible linker into the first multiple cloning site of pCDF- Duet vectors retaining the NdeI and HindIII restriction sites. Sequence-verified constructs were transformed into T7 Express *E. coli* cells.

### Purification of amidases P203, P204 and P205

Production of P203, P204, P205 was performed as previously described.^26^ Solution used for protein purification are described in **Table S2**. In short, *E. coli* BL21 was induced in exponential phase (OD ∼ 0.5) using 0.1 mM isopropyl β-D-1-thiogalactopyranoside and grown for 2 days at 16°C and 250 rpm with 50 µg/mL spectinomycin. The cell cultures (50 mL) were pelleted, resuspended in lysis buffer and sonicated. The lysate was loaded onto a pre-equilibrated column packed with Ni-NTA agarose beads (Qiagen), subsequently washed twice with wash buffer, and the his-tagged amidases eluted with elution buffer. The purified protein fractions were desalted using a PD-10 column (Cytiva) and eluted in SGT buffer. The protein concentration of the eluate was determined using the Coomassie (Bradford) protein assay (Thermo Fisher Scientific) with comparison to a bovine serum albumin standard curve. The protein eluate purity and size was assessed using SDS-PAGE with Mini-Protean TGX Precast Gels (Bio-Rad) and the Page Ruler Plus Prestained Protein Ladder (Thermo Fisher Scientific).

### Catalytic inactivation of P205 by site-directed mutagenesis

To create a catalytically inactive variant of the paracetamol amidase homolog P205, serine 146 in the catalytic triad (Ser146-Ser170-Lys71) was substituted with alanine based on Shin et al. 2003.^27^ Site-directed mutagenesis introduced point mutations at residue 146 (GCT instead of TCT) using primers Lb AS S146A fw (CAG TCC TGG TGG GGC TTC GG) and Lb AS S146A rev (TAG GCC AGG TTC CAG GGA TTG CG). Mutagenesis involved PCR amplification with Q5 high fidelity Mastermix (New England Biolags), kinase-ligase-DpnI (New England Biolabs) ligation of vector and removal of template DNA, and transformation into *E. coli* DH5α (New England Biolabs), followed by plasmid isolation (Qiagen Plasmid Miniprep), and sequencing to confirm the correct clones by Sanger sequencing (Microsynth, Switzerland)

### Paracetamol in vitro biotransformation by P205

A solution of 50 mg/L paracetamol in 50 mM Tris-HCl (pH 8) was mixed with purified P205 negative control. The positive control was paracetamol amidase ANS81375.1 (0.087 mg/mL). Reactions were incubated at 37°C and 225 rpm, with samples taken at various times, stopped with acetonitrile, and centrifuged. Supernatants were diluted 1:10 with MilliQ water and stored at 4 °C until HPLC analysis. 20 μL samples were separated using a Nucleosil RP 18 HD column (Macherey-Nagel) with a gradient elution of buffer A (20 mM phosphate buffer, 0.04% phosphoric acid, pH 3.0) and buffer B (90% acetonitrile, 0.04% phosphoric acid, pH 3.0). The elution gradient was set as follows: 2% – 100% buffer B over 10 minutes, followed by 5 minutes isocratic elution at 2% buffer B, at a flow rate of 0.8 mL/min. Paracetamol was detected at 250 nm with a retention time of 6.6 minutes.

### Biochemical characterization of P205

Biochemical characterization of P205 was performed using established colorimetric protocols for esterase activity.^28^ In short, to determine the pH optimum of P205, the standard esterase substrate 4NP trimethylacetate was used, adjusting for background hydrolysis rates at different pH levels. Measurements of 4-nitrophenol (4NP) were taken at its isosbestic point wavelength of 347 nm for robust pH assay results, as demonstrated by Peng et al.^29^ In a 96-well plate, 200-µL reactions were prepared using 100 mM citrate buffer at pH 4, 5, and 6, and 100 mM Tris-HCl buffer at pH 7, 8, and 8.5, with measurements taken every 2 minutes. The substrate 4NP trimethylacetate was used and standard curves of 4NP generated for each pH. To determine the temperature optimum of P205, absorbance of 4NP at 410 nm in Tris-HCl buffer (pH 8) was measured at different temperatures using 4NP-trimethylacetate as substrate. The thermal stability of p205 was assessed by measuring 4NP absorbance at 410 nm in Tris-HCl buffer (pH 8) after incubating the enzyme at various temperatures (37°C, 39.5°C, 51.5°C, 60.5°C, 70.0°C, 81.2°C, 90.0°C) for 1 hour using 44NP-butyrate as substrate. All experiments were conducted using purified P205 at a concentration of 0.0022 mg/mL and the catalytically inactivated variant, P205-S146A, was used as negative control.

### Substrate specificity screening of P205 using micropollutants

Substrate specificity screening of P205 on a collection of 183 micropollutants, including commonly used pesticides, artificial sweeteners, and pharmaceuticals, was done as previously described.^30^ A detailed description is available in the Supplementary Information. In short, a total of 183 micropollutants were divided into 14 submixtures and added to a 96-well plate, achieving a final working concentration of 200 μM for each micropollutant in 100 mM Tris-HCl buffer (pH 8). Purified P205 enzyme was added to reach a final concentration of ∼190 nM. P205-S146A served as a negative control. Samples were collected at 0h, 6h, and 24h, and the reactions were quenched using methanol. After centrifugation, the supernatant was collected and stored for subsequent measurement using ultrahigh-performance liquid chromatography coupled with high-resolution tandem mass spectrometry (UHPLC-HRMS/MS). Details on LC-HRMS can be found in the Supplementary Information.

### Bioinformatic analysis of amidase signature family enzyme sequences

The genomes of microbial isolates from the catheterized urine of female patients previously described by Thomas-White et al. were used in this analysis.^11^ Of the 149 bacterial isolates retrieved,11 129 genome assemblies with matching strain classifications could be retrieved from NCBI and were used in the study. Sequence alignments of the amidase P205 and ANS81375.1 sequence^16^ to reference genomes were achieved by a reverse protein alignment search with BLAST+.^31^ Significance cutoffs of e-value ≤ 0.1, query coverage ≥ 20 % and bit-score ≥ 50 were used. IQ-TREE was used to estimate the phylogenetic relationship of the 157 protein sequences.^32^ A maximum-likelihood phylogenetic tree of paracetamol amidase homologs of urinary bacteria was constructed using IQ-TREE with amino acid substitution model LG+F+I+G4 based on 1000 ultrafast bootstraps.^33,34^ The function of the amidase homologs was predicted using a contrastive learning method for enzyme function prediction (CLEAN).^20^ If multiple EC numbers were predicted, only the highest confidence EC is shown in the figure. A subsequent maximum-likelihood phylogenetic tree of 16 active amidases was constructed using IQ-TREE with amino acid substitution model LG+I+G4 based on 1000 ultrafast bootstraps.^33,34^

### Amidase signature family enzyme library construction

The 157 unique proteins were clustered at a 60% amino acid identity threshold using CD-HIT v4.8.1 with default parameters.^35^ Sequences with over 90% amino acid identity to the query sequence and sequences without a ‘GSS’ motif characteristic of the amidase signature family^18,36^ were excluded. Cluster sizes were reduced using the Hamming distance to maximize sequence diversity while retaining a maximum of 50% of the sequences per cluster. Within each cluster, for sequences originating from the same genus, only the sequence with the highest percentage identity to the query sequence was retained. We further removed all predicted EC 6.3.5.7 sequences (glutaminyl-tRNA synthases) and kept a maximum of three sequences for a given genus (retaining those with highest similarity to the query sequence). We further removed sequences that were similar to previously characterized sequences using PaperBLAST^37^ e.g., such as previously-characterized colibactin amidase of *E. coli*,^38^ to obtain a final set of 37 new predicted aryl acyl-amidase proteins to test for activity (**Table S3**).

### Substrate specificity screening of urinary microbiota amidase library

Based on the results obtained from the substrate specificity screening with P205, we observed that accepted substrates had aryl amide substructures. However, several other amides in the xenobiotic library (e.g., rufinamide, mirabegron, benserazide) were not hydrolyzed. To understand which structural/chemical properties of the substrate influence the activity of the amidases, we selected an additional set of 17 low-molecular weight organic compounds with different aromatic ring substituents and hydrolyzable moieties (**Figure S1**).

### HPLC-based enzyme assays

To screen 17 organic compounds for biotransformation with 40 candidate aryl acylamidases (n = 680 candidate enzyme-substrate combinations), the substrates were pooled to control the number of samples and enable experiments to be run in triFplicates. Pool 1 consisted of acesulfame, atenolol, capecitabine, chlorpropham, diuron, vorinostat, and pool 2 consisted of acetylsulfamethoxazole, atorvastatin, dasatinib, metoclopramide, propanil, rufinamide. Stock solutions of individual compounds were prepared in DMSO at a concentration of 10 mM. The final working concentration for each compound was 100 µM (see **Table S1** for supplier list).

Induced cells (1 mL volumes) were lysed using the BugBuster® Protein Extraction Reagent (Sigma-Aldrich) according to the manufacturer’s instructions without use of any optional steps. The obtained lysate was then diluted with 250 μL Tris-Hcl pH 8. 150 μL lysate was added to 1350 μL 50 mM Tris-HCl pH 8 containing pool compounds at final concentration of 100 μM each in a 2mL Eppendorf tube. The tubes were incubated at 37°C with no agitation for 24 h. Timepoint t0 and t24 were taken by mixing 500 μL samples and stopping the reaction by addition of 500 μL acetonitrile. The obtained samples were run on the HPLC.

Reaction substrates and products were quantified using HPLC coupled to a UV-vis detection unit (Dionex Ultimate 3000 Systems, Thermo Scientific). Twenty microliters sample were injected from an autosampler cooled to 10°C into a Nucleoshell RP 18plus column (particle size 5 μm, 150 x 3 mm, Macherey-Nagel) and eluted with 20 mM phosphate, 0.04% phosphoric acid (A), and 90% acetonitrile, 0.04% phosphoric acid (B) at a flow rate of 0.8 mL/min. The elution gradient was set as follows: 100% A: 0 – 5 min, 0% – 80% B: 5 – 25 min, and 100% A: 25 – 30 min. The UV-Vis detector measured at wavelengths 210, 225, 252 and 305 nm. Retention times for the first pool of substrates were as follows: acesulfame (3.8 min), atenolol (9.2 min), capecitabine (14.9 min), chlorpropham (21.3 min), diuron (17.7 min), and vorinostat (13.4 min). For the second pool of substrates, the retention times were: acetylsulfamethoxazole (14.0 min), atorvastatin (21.2 min), dasatinib (14.8 min), metoclopramide (11.8 min), propanil (19.2 min), and rufinamide (13.0 min).

### Colorimetric enzyme assays

Biotransformation of 4NP-butyrate, 4NP-trimethylacetate, flutamide, and nitroacetanilide was performed in diluted crude lysate (using the BugBuster® lysis procedure). The procedure is adapted from established colorimetric assay as previously described.^28^ Briefly, 20 μL of crude lysate was added to 180 μL of 50 mM Tris-HCl (pH 8) buffer containing 200 μM substrate in a 96-well microtiter plate. The P205-S146A mutant was used as a negative control. A solution of 200 μM nitrophenol, 4-nitro-3-(trifluoromethyl)aniline, and nitroaniline in 50mM Tris-HCl (pH 8) served as complete substrate turnover reference. The plates were incubated at 37°C in a BioTek Synergy H1 (Agilent) plate reader with linear shaking, and the optical density (OD) was measured at 410 nm. The incubation times were approximately 12 hours for 4NP-butyrate and 4NP-trimethylacetate, and approximately 24 hours for flutamide and nitroacetanilide.

For paracetamol, a whole-cell biotransformation assay was performed. Single colonies of each member of the amidase expression library were grown at 37°C in LB with 50 μg/mL spectinomycin and induced in exponential phase with 0.1 mM IPTG, followed by overnight incubation at 25°C. Each well in a 96-well microtiter plate was filled with 180 μL of PBS containing 10% LB, 50 μg/mL spectinomycin, and 500 μM paracetamol. Subsequently, 20 μL of the induced bacterial culture was added to each well in technical duplicates. *E. coli* T7 Express P205 was used as the positive control, and *E. coli* T7 Express P205-S146A served as the negative control. The plates were incubated at 37°C in a BioTek Synergy H1 (Agilent) plate reader with linear shaking, and the OD was measured at 400 nm over approximately 24 hours. Additionally, a standard curve with the paracetamol hydrolysis product, 4-aminophenol, ranging from 4 mM to 62.5 μM, was prepared in PBS with 10% LB and 50 μg/mL spectinomycin.

Biotransformation of substrates 4NP-butyrate, 4NP-trimethylacetate, flutamide, nitroacetanilide, and paracetamol was assessed by measuring the absorbance of the transformation products. Specifically, the chromophore 4NP was measured at 410 nm for both 4NP-butyrate and 4NP-trimethylacetate, 4-nitro-3-(trifluoromethyl)aniline at 410 nm for flutamide, 4-nitroaniline at 410 nm for nitroacetanilide, and oxidation products of 4-aminophenol at 400 nm for paracetamol.^39^

### Data analysis

The HPLC and plate reader data were processed using R v4.4.0 and RStudio v2024.04.2+764. Chromeleon 7.2.10 ES (Thermo Scientific) was used to integrate the peaks and obtain the peak absorption areas of samples measured by HPLC. Compound area chosen based on best absorption at the measured wavelengths 225 nm (acesulfame, atenolol, chlorpropham, rufinamide), 252 nm (propanil, diuron, atorvastatin, acetylsulfamethoxazole, vorinostat) and interquartile range (IQR) for P205-S146A were removed. The relative removal of each sample was calculated using the following formula:

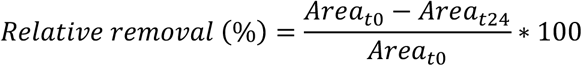

For hit-calling of substrates analyzed by HPLC two conditions needed to be met: (1) median removal higher than the median + 1.5 x IQR of P205-S146A, and (2) relative removal over 50%. The hit calling for substrates analyzed by the plate reader was conducted at specific times depending on the substrate: 100 minutes for 4NP-butyrate, 600 minutes for 4NP-trimethylacetate, 1250 minutes for nitroacetanilide, 1250 minutes for flutamide, and 2600 minutes for paracetamol. Hits were identified if the mean yield (OD at specified time) was at least 2 standard deviations higher than that of P205-S146A, and for 4NP-butyrate and 4NP-trimethylacetate, the yield had to be over 30%, with yield calculated as:

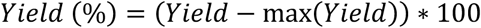

The amino acid sequences of the 16 enzyme hits were queried in the Global Microbial Gene Catalog (GMGC) in September 2023 using the minimal homology threshold. AlphaFold3 models^40^ of the 16 enzyme hits were generated using the AlphaFold Server in May and June 2024. AlphaFold3 structures were visualized in PyMol (v2.5.4).

### Machine learning

For the alignment of amidase amino acid sequences, we trimmed the alignment to include only the amidase signature region comprising approximately 160 amino acids.^21^ To featurize the amino acids, we used the 4 first principal components corresponding approximately to polarity, secondary structure, molecular size or volume, and codon diversity as described in Atchley et al.^41^ To featurize the chemicals, SDF files were generated from isomeric SMILES, and the `atomcountMA` function from the ChemmineR package was used to calculate the frequency of atoms and functional groups. Additionally, the following features were manually engineered (in parentheses the name used in the text): ANILIDE (aryl amide), MW_bond (amide tail mass, representing the molecular weight of the amide carbonyl substituents), MW_ring (aryl mass, molecular weight of the aryl ring with all substituents), and MW (molecule mass, molecular weight of the molecule). Special cases for non-anilides included acesulfame (MW_ring and MW_bond are each half of the MW), atenolol and rufinamide (MW_bond is a primary amine hydrogen, MW_ring includes the methyl group attached to the amide for atenolol), esters (4NP-butyrate and 4NP-trimethylacetate, were treated as if the ester was an amide moiety), and metoclopramide (treated as if it was an anilide). Low variance features were removed using the `nozerovar` function. Data were split using random sampling into training and testing sets, stratified according to enzyme-substrate activity (1: active, 0: inactive), with the training set consisting of 80% and the testing set 20% of the data. Ten individual XGBoost models were trained, and a final model was trained using features present in at least 70% of the individual models to assess feature importance.

## Results and discussion

### Biochemical characterization of the aryl acylamidase P205

Previously, we found that genomes from the urinary microbiota encode sequence homologs of enzymes involved in various micropollutant biotransformations.^15^ In particular, we identified the widespread conservation of homologs of paracetamol amidases^15^ belonging to the AS enzyme family in urinary bacteria. Here, we selected three different AS enzymes (here denoted P203, P204 and P205) from three different urinary bacterial taxa (*Corynebacterium, Gordonia,* and *Lacticaseibacillus*, respectively) and cloned the respective genes into inducible expression vectors in *E. coli*. Among these, P203 and P204 did not express in *E. coli,* however the gene encoding P205 from *Lacticaseibacillus rhamnosus* expressed well and could be purified to homogeneity (**Figure S2**). We biochemically characterized P205 and measured an enzymatic activity temperature optimum of 40°C, a pH optimum of 8, and heat stability up to 60°C (**Figure S3**). The temperature and pH optima of P205 were similar to previously reported AS enzymes AmpA^42^ and TccA^43^, but P205 retained almost complete activity after 1 hour at 60°C, unlike the significant activity reduction observed in AmpA and TccA. We next incubated purified P205 with paracetamol and observed paracetamol amidase activity (**Figure 1A**) confirmed by UHPLC-HRMS/MS (**Figure 1B**). To assess the substrate range beyond paracetamol, we used an established 96-well UHPLC-HRMS/MS screening assay^30^ to test P205 for activity with 183 wastewater-relevant micropollutants. In order to account for potential non-specific effects such as substrate sorption, we generated a catalytically inactivated enzyme variant of P205 by exchanging the catalytic serine 146 with alanine. This method for catalytic inactivation was reported previously for AS enzymes^23,27^ and we confirmed here that it yielded an inactive enzyme. Through comparison to our inactivated enzyme control, we detected biotransformation (>40% removal of substrate) by P205 for two pharmaceuticals in addition to paracetamol: vorinostat and capecitabine (**Figure 1B**). The stringent removal threshold was

**Figure 1:**
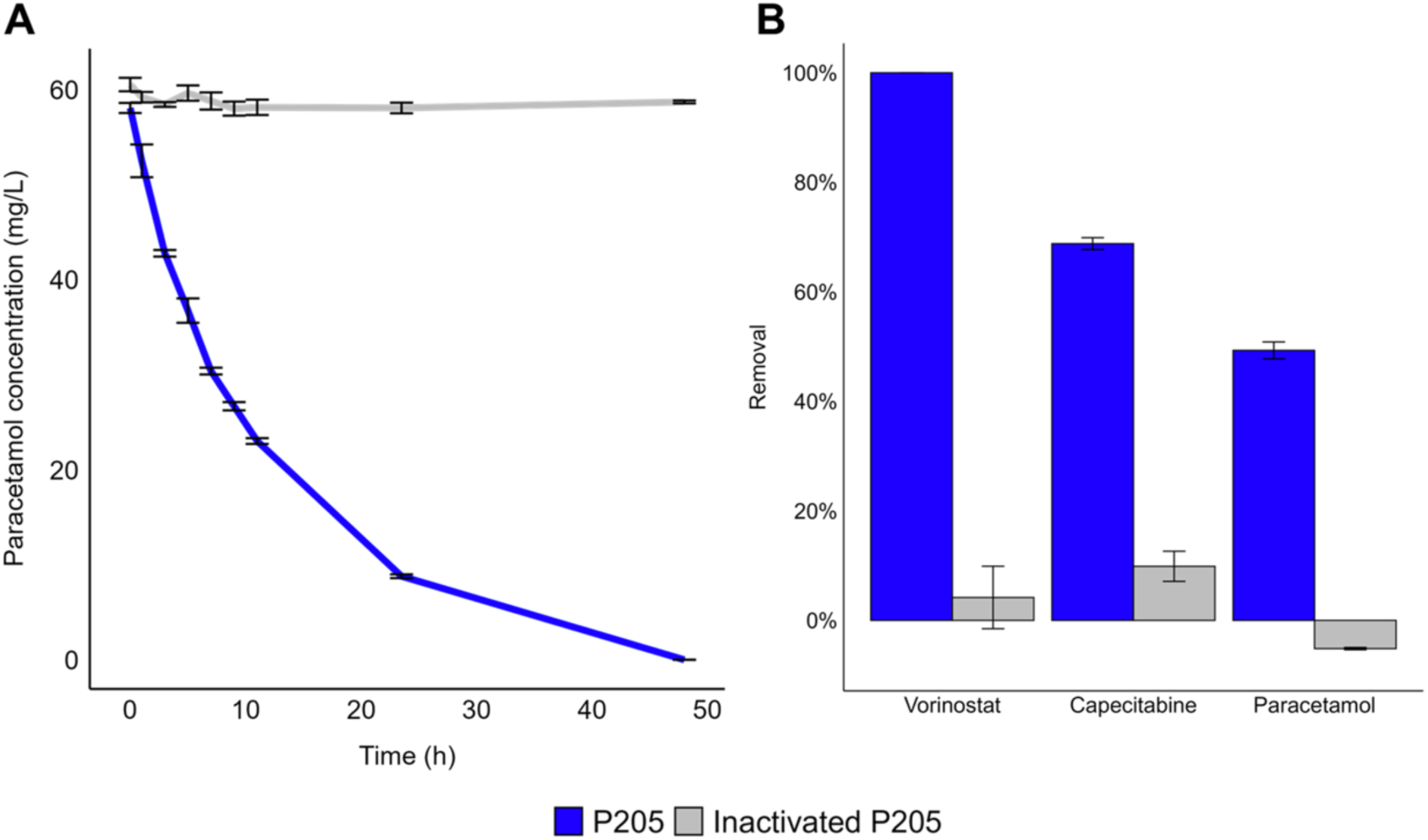
**A)** Paracetamol removal curve by P205 (blue). The inactive enzyme control (gray) is the catalytically inactivated variant P205-S146A. **B)** Substrate specificity screening hits after 24 h incubation. chosen to exclude false positives, such as compounds without hydrolyzable moieties or those with high removal variability (**Figure S4**). Based on the results obtained from the substrate specificity screening with P205, we observed that accepted substrates had aryl amide substructures. However, several other amides in the xenobiotic library (e.g., rufinamide, mirabegron, benserazide) were not hydrolyzed. We additionally confirmed amide bond cleavage of capecitabine by HPLC using 5’-deoxy-5-flourocytidine as an authentic standard (**Figure S5**). Overall, based on these results, we identified aryl amide moieties to be a common feature of the three P205 substrates.

### Urine-derived microbial amidase library selection

These findings prompted us to investigate whether other urine-derived microbial AS enzymes exhibited an expanded substrate specificity beyond paracetamol including different pharmaceuticals and other micropollutants. To systematically sample AS enzyme diversity, we expanded our analysis to a larger panel of bacterial isolates from urine including isolates from patients with diagnosed urinary tract infections,^11^ whose urinary microbiota might be exposed to higher pharmaceutical loads. To select candidate enzymes we used the active AS enzyme from *L. rhamnosus* P205 as a query sequence (see Methods for more detail on the selection procedure). A homolog of P205 was found in 123 out of 128 (96%) genomes queried.

A phylogenetic tree of the unique hits (n=157) revealed two main clades (**Figure 2**) which correspond to two different predicted enzyme functions.^20^ Over half of AS enzyme homologs were predicted to be glutaminyl-tRNA synthases (EC 6.3.5.7)^21^ while 40% were predicted to be C-N hydrolases (EC 3.5). Given our focus on amide bond hydrolysis, we selected and synthesized 37 of the predicted C-N hydrolases to construct a diverse AS enzyme library spanning 22 different bacterial genera.

**Figure 2:**
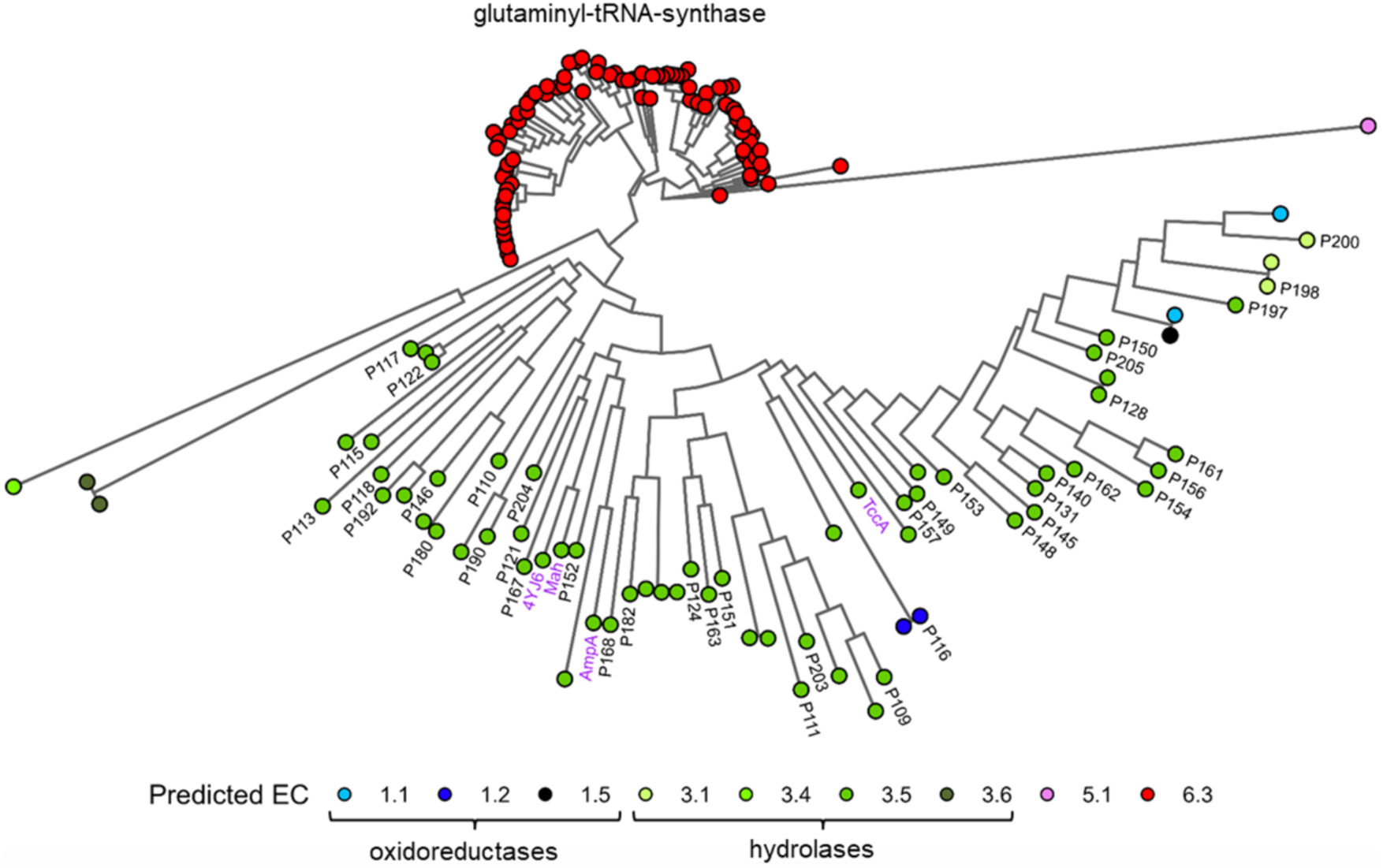
Phylogenetic tree of AS enzyme homologs from the urinary microbiome. Circles are colored by predicted EC number. Black labels: homologs selected for the AS enzyme library.

### Urine-derived microbial AS enzymes have variable substrate specificities

To profile substrate specificity of the urinary AS enzyme library (n=40 enzymes), we selected a set of 17 structurally-diverse compounds with hydrolyzable moieties, primarily amides and esters including a subset of the 183 wastewater relevant micropollutants previously tested with P205 (see Methods for selection criteria). We expressed each AS enzyme in our library (including P203, P204, P205) individually in *E. coli* strains and prepared crude lysate extracts for high-throughput substrate screening^30^ for a total of 680 distinct enzyme-substrate combinations.

Of the 40 AS enzymes tested, 16 enzymes were active on at least one substrate and are further analyzed here (**Figure 3**). The most active AS enzymes in our library were the phylogenetically-related P131 and P148 from *Streptococcus salivarius* and *Aerococcus sanguinicola*, respectively, which transformed 9 out of 17 substrates tested (**Figure 3A and 3B**). In comparison to the rest of the AS enzyme library, P205 showed weak activity in crude lysate (**Figure S6 and S7**). It did not meet the stringent activity thresholds (see Methods) set to limit false positives for downstream analysis and modeling. Apart from the 4-nitrophenyl (4NP) compounds included as standard hydrolase substrates, the micropollutants biotransformed by the highest number of enzymes were paracetamol and the herbicide propanil (**Figure 3B**). Activity on these compounds has also been reported by other microbial AS arylamidases.^16,22,42–44^ In contrast to the previously characterized enzymes, AS enzymes in the library did not biotransform chlorpropham. Additional compounds not transformed by any AS enzymes in the library included acesulfame, atenolol, atorvastatin, dasatinib, diuron, metoclopramide and rufinamide. In addition to known substrates, we also discovered new AS enzyme family substrates. To our knowledge, no AS family enzymes have previously been reported to biotransform vorinostat, capecitabine, flutamide or acetylsulfamethoxazole.

**Figure 3:**
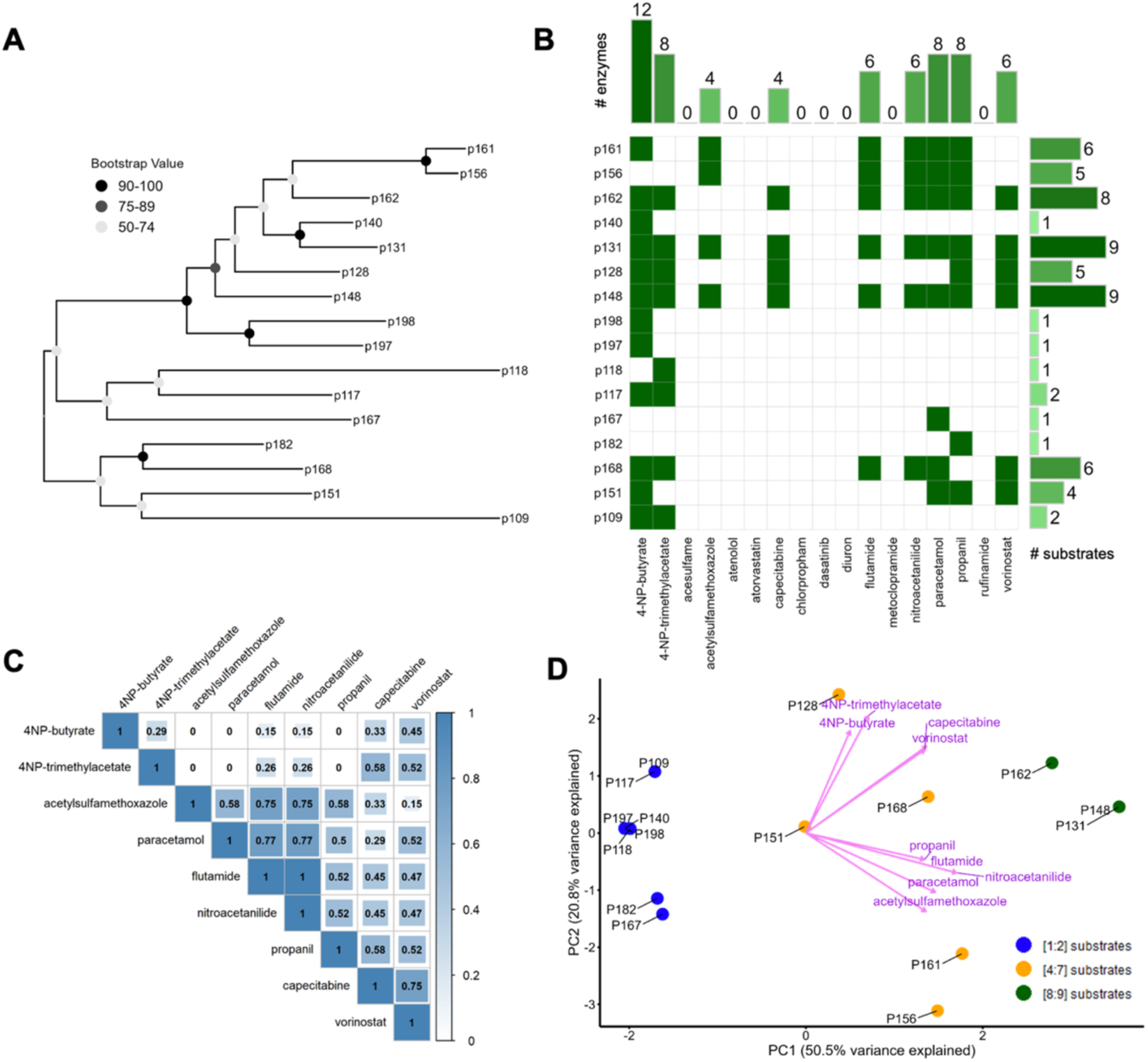
**A)** Maximum-likelihood phylogenetic tree of 16 microbial urine-derived AS enzymes active on at least one tested substrate. Bootstrap confidence levels are shown in grayscale from 1000 bootstrap replicates. **B)** Heatmap of enzyme-substrate hits. Barplots show the number of enzymes active on a given compound (x-axis) and number of substrates of a given enzyme (y-axis). **C)** Pearson correlation plot of substrates accepted by at least one enzyme. **D)** PCA biplot of enzyme-substrate matrix. Pink arrows visualize the principal component 1 and 2 loadings and group roughly based on conserved chemical moieties e.g., 4-nitrophenyl esters, short and long aliphatic chains. Points are colored by the number of accepted substrates of the enzymes.

A substrate specificity dendrogram, based on the Jaccard similarity index (excluding substrates not biotransformed by any enzyme), revealed three distinct clades for AS enzymes that are active on at least four substrates (**Figure S8**). Interestingly, not only phylogenetically-related enzymes like P156 and P161 share similar substrate specificities, but also distantly related enzymes P168 and P162. Substrate specificity only partially correlates with protein phylogeny, suggesting convergent evolution of biotransformation capabilities. The average amino acid identity between all of the 16 active enzymes is only 29% (± 8%) (**Figure S9**), indicating that AS enzymes with diverse sequences are also capable of catalyzing similar reactions.

A Pearson correlation matrix of substrates accepted by at least one enzyme (**Figure 3C**) reveals correlated substrate preferences for e.g., low molecular weight substrates (flutamide, nitroacetanilide, paracetamol) and substrates with long aliphatic tails (capecitabine, vorinostat). Principal component (PC) analysis of the accepted substrates (**Figure 3D**) shows enzymes cluster by their number of accepted substrates, especially along PC1. While acknowledging the limitations of PC analysis,^45^ this suggests that key chemical moieties in accepted substrates (e.g., aliphatic chain length, type of hydrolyzable bond) may be used to predict the degree of promiscuity of AS enzymes.

### Machine learning model identifies 24 features predictive of AS enzyme-substrate relationships

To gain additional insights into the physicochemical features underlying enzyme-substrate specificity, we trained machine learning models on enzymes active with at least one substrate (n=272 enzyme-substrate combinations). Among the models tested, we selected an extreme gradient boosting decision tree algorithm (XGBoost) for feature selection of physicochemical properties from enzymes and substrates (see Methods).

We trained a final XGBoost model using top selected enzyme and chemical features and achieved an F1 score of 0.94, balanced test set classification accuracy of 89%, sensitivity of 93% and specificity of 85%. The receiver operator characteristic (ROC) curve shows an AUC of 0.912 (95% confidence interval: 0.809 - 1.0) (**Figure S10**). The top ten features included a balanced mixture of chemical and enzyme features, suggesting their combination played a role in enzyme-substrate specificity (**Figure 4**). As the model was limited to the narrow set of substrates and AS enzymes included in our library, our goal was not to optimize model performance but rather to extract and interpret fine-grained features driving enzyme-substrate specificity in our dataset. While acknowledging the constraints, we viewed this as an informative pilot study towards training more generalizable models to predict enzymatic micropollutant biotransformations across enzyme classes.

**Figure 4:**
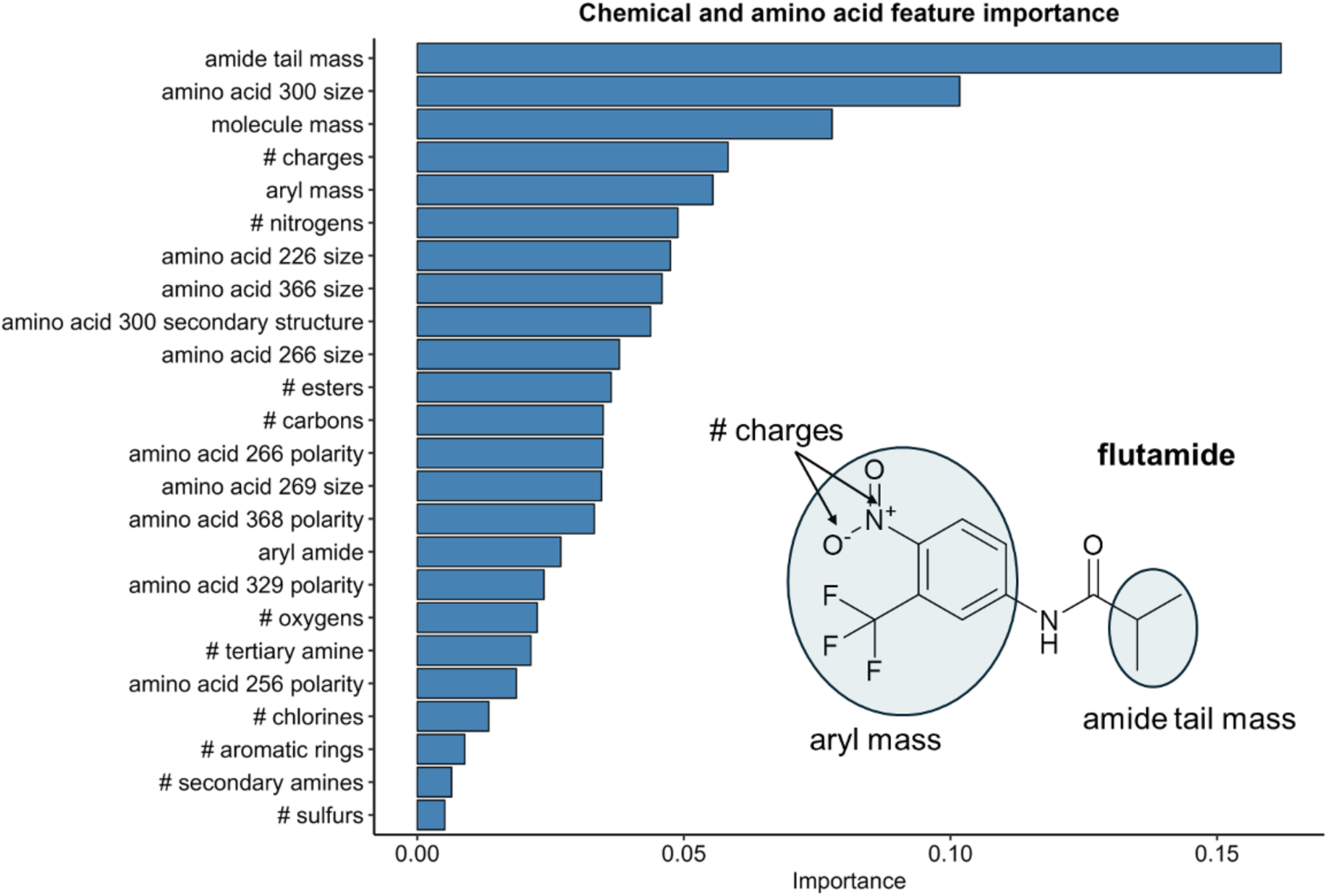
Chemical and amino acid feature importance in a XGBoost model trained using the most features derived from ten individual models. Amino acid numbers refer to positions in the protein alignment of 16 AS enzyme hits. The chemical structure inset depicts two custom labels used for chemical featurization which were found to have high feature importance weighting (see Methods for details on chemical featurization).

### Amide tail and aryl substituent charges influence enzymatic activity

The molecular weight of the amide carbonyl group substituent (referred to here as the ‘amide tail’) was the most important feature that was predictive of enzyme-substrate activity (**Figure 4**). Indeed, the amide-containing compounds which were the most widely biotransformed (paracetamol, propanil, nitroacetanilide, flutamide) contained smaller amide carbonyl group substituents compared to compounds which were not biotransformed (e.g., atorvastatin and dasatinib). Vorinostat and capecitabine were still biotransformed suggesting that even large substituents were accepted, with an apparent preference for linear saturated aliphatic chains. In comparison with X-ray crystallography data from a bacterial acyl arylamidase co-crystallized with its substrate paracetamol (Protein Data Bank ID: 4YJI), the substrate was co-crystallized with the amide carbonyl substituent oriented towards the interior of the binding pocket.^23^ The authors identified a substrate tunnel capable of accommodating compounds with *N*-acyl groups up to C_10_ in length^23^ (similar carbon chain lengths to vorinostat and capecitabine). Higher molecular weight substituents, such as those found in dasatinib and atorvastatin, were not accepted by AS enzymes, suggesting that these may not fit into the substrate tunnel.

The molecular weight of the substrates was also a predictive feature consistent with the relatively low molecular weight of most preferred AS enzyme substrates including paracetamol and propanil. The formal charges of several favored substrates including 4NP-butyrate, 4NP-trimethylacetate, flutamide, and nitroacetanilide was also an important feature. These compounds all bear electron-withdrawing nitro groups in the *para*-position on their phenyl rings, increasing susceptibility to nucleophilic attack at the carbonyl carbon. For example, Ko et al. observed four orders of magnitudes higher V_max_/K*_M_* for the reaction with 4-nitroacetanilide compared to acetanilide with an AS family enzyme.^22^ The susceptibility of 4-nitrophenyl compounds to hydrolysis is evident in their use as activity probes for various enzyme families.^28,46–49^

### Identified residues are located on the enzyme surface

The size and secondary structure of amino acid position 300 in the alignment of the 16 active amidases was important for AS enzyme-substrate specificity (**Figure 5A**). Other amino acid features relevant for model prediction included the size and polarity of amino acid 226, the size of amino acid 266, and the size of amino acid 366. To identify whether certain amino acids were associated with the number of substrates accepted, we classified the substrate specificity of our AS enzymes as ‘promiscuous’ (eight or more substrates) or ‘narrow’ (two or fewer substrates). We then visualized the amino acid sequence logos for ‘promiscuous’ (P131, P148 and P162) and ‘narrow’ enzymes (P109, P117, P118, P140, P167, P182, P197 and P198) of the eight alignment positions identified by our gradient boosting model to be important (**Figure 5B**). Apart from position 269 where asparagine (N) is present solely in all promiscuous enzymes, other positions are highly variable, although there is only a small overlap between amino acids of promiscuous versus non-promiscuous enzymes (**Figure 5A**). This emphasized the limitations of our model for interpreting some identified features which do not have a readily apparent biological or chemical role yet correlated well with enzyme-substrate patterns.

**Figure 5:**
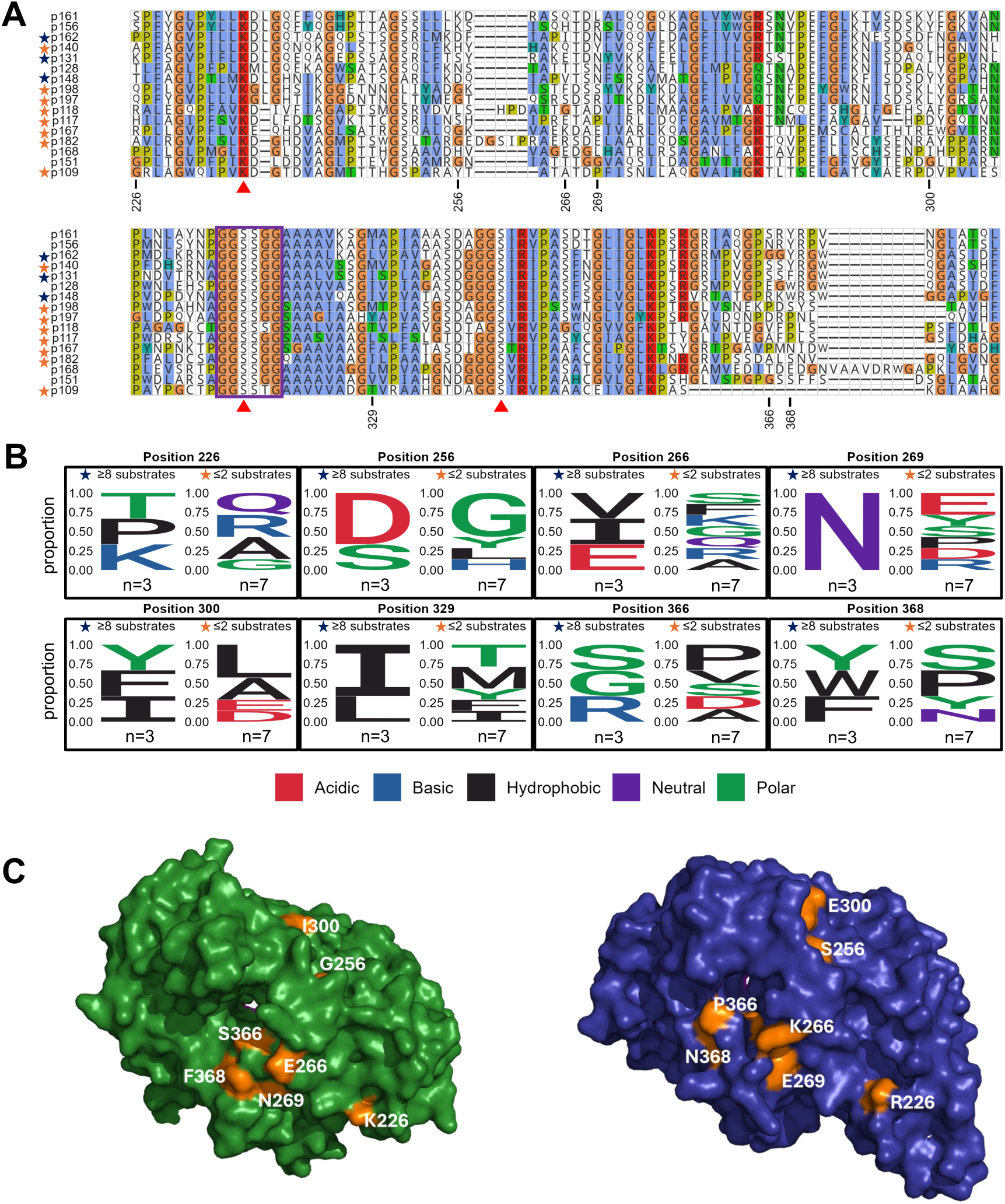
**A)** Multiple sequence alignment of the amidase signature region of library enzymes active with at least one substrate. The sequences are ordered based on the phylogenetic tree shown in Figure 3. The numbered positions correspond to those with high feature importance scores shown by XGBoost. Red triangles mark the catalytic triad residues and the violet box marks the Gly/Ser-rich motif. Black stars indicate ‘promiscuous’ enzymes and orange stars are for ‘narrow’ substrate range enzymes. **B)** Sequence logos of amino acids positions (in the alignment) with high feature importance by XGBoost. **C)** AlphaFold3 structures of P131 (green) and P167 (blue) with view on the pocket entrance. Orange: residues corresponding to those with high feature importance scores shown by XGBoost. Residue numbers refer to positions within the Figure 3 sequence alignment.

Next, we visualized important amino acids in AlphaFold3 protein structural models to identify their localization and infer potential biochemical relevance (**Figure 5C**). With the exception of amino acid 329, all residues were situated on the surface of the proteins. In order to identify the predicted substrate tunnels, we structurally aligned AlphaFold3 models of the 16 active amidases (average RMSD 2.275 ± 1.901 Å) with 4YJI. Residues 266, 269, 366, and 368 mapped near the probable substrate tunnel entrance.

Previously, Lee et al. (2015) identified residues forming a hydrophobic cavity around the acyl group of paracetamol to the substrate through hydrophobic interactions. They localized residues influencing substrate binding and stabilization through hydrogen bonds.^23^ Zhang et al. (2023) additionally identified Tyr138 as an important residue in the binding pocket that determines substrate specificity.^44^ These previous studies did not investigate residues on the surface of the protein. Our investigation identified surface residues as additional important positions which correlated with AS enzyme-substrate specificity and promiscuity. Further experimental validation e.g., through site-directed or alanine scanning mutagenesis, will enable validation of the importance of these residues.

### Environmental relevance

Having established the promiscuity of AS enzymes in micropollutant biotransformations, we next investigated how widespread and conserved AS enzymes were across different microbiomes. We queried the amino acid sequences of our 16 AS enzymes in the Global Microbial Gene Catalog (GMGC).^50^ The majority of homologs were detected in human-associated samples (including gut, oral, skin) reflecting the urinary sampling locations of our sequences and overall greater sequencing efforts towards human-associated habitats (**Figure S11A**). About 20% of homologs are represented in the environmental microbiomes built environment, soil, and wastewater. When only considering these ecosystems, most enzyme homologs were found in soil and the built-environment (**Figure 6**).

**Figure 6:**
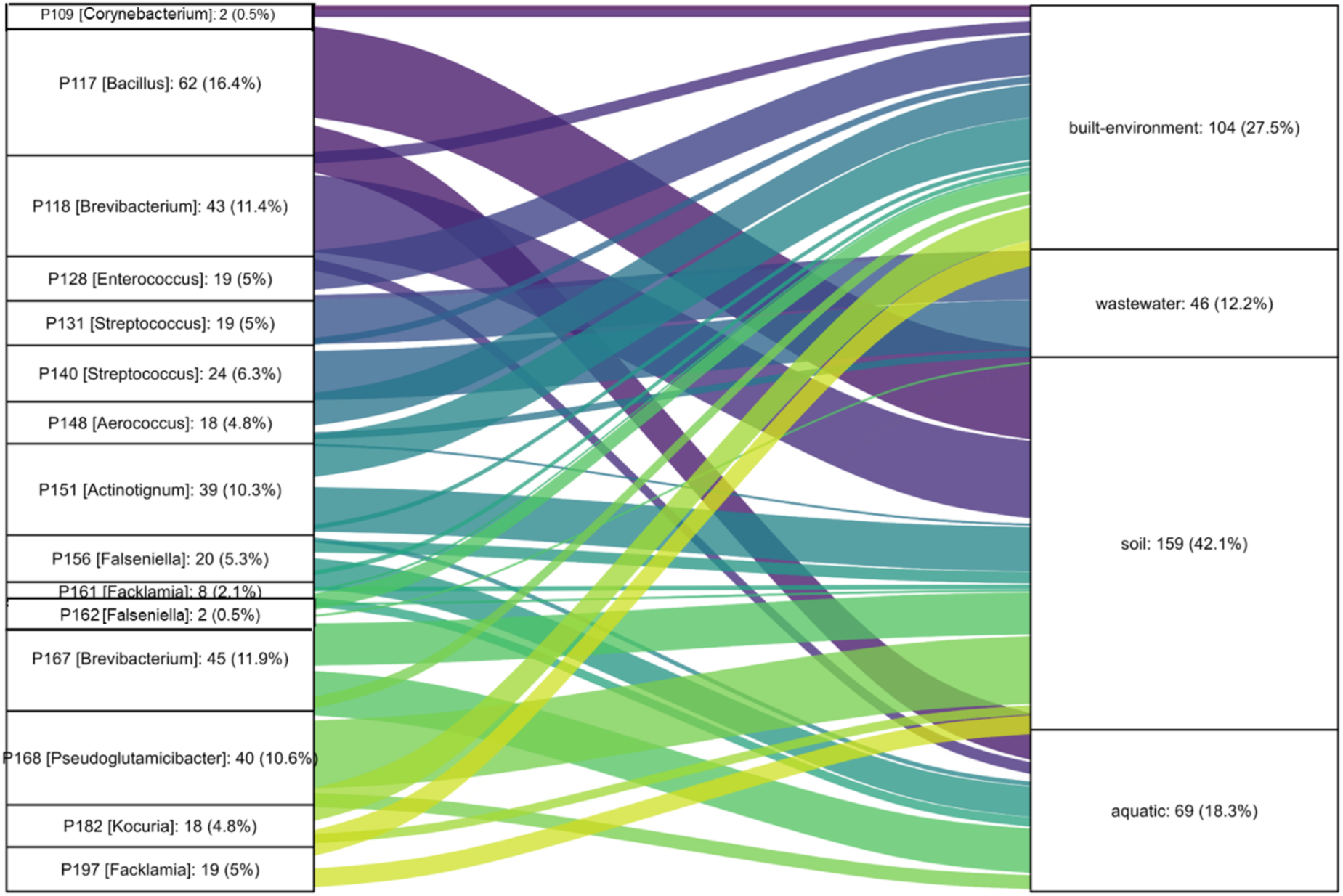
Distribution of enzyme homologs across environmental microbiomes (according to the Global Microbial Gene Catalog^50^). Each stratum represents an enzyme, with the bacterial host genus indicated in brackets. The connections between strata represent the prevalence of each enzyme in various environments. The labels on the strata display both the absolute counts and the relative percentages of the enzyme homologs. For clarity, only connections with values greater than 3 are shown.

Examining the most promiscuous enzymes (P131 and P148) in our dataset, we detected P131 (*Streptococcus*) homologs in wastewater, while P148 (*Aerococcus*) homologs were predominantly detected in the built environment and soil (**Figure S11B**). The global abundance distributions of both enzymes were nonetheless dominated by human-associated microbiomes. Their detection in human-associated contexts warrants future research on the potential of AS-mediated microbial biotransformations for modulation of anticancer treatment efficacy,^51^ since P131 and P148 readily biotransformed the anticancer drugs flutamide, capecitabine, and vorinostat. Also relevant for human-associated contexts, these and six additional AS enzymes also biotransformed the widely-used analgesic, paracetamol.

Paracetamol was among the most preferred aryl amide substrates across all AS enzymes measured in our study. In support of this finding, diverse AS enzymes were previously shown to be involved in the biotransformation of paracetamol in wastewater^52^. Despite the ready biodegradability of paracetamol, this parent compound is still often measured in µg/L in concentrations in both wastewater effluent^53^ and natural surface waters.^54^ A 2022 global river survey identified paracetamol as the most abundant compound (average 3.1 µg/L) out of 61 different pharmaceuticals measured.^54^

Incomplete paracetamol removal during wastewater treatment also has ‘downstream’ effects as was recently demonstrated in agricultural irrigation systems using recycled wastewater effluent.^55^ Strikingly, paracetamol concentrations in the low µg/L range in wastewater effluent used for irrigation induced shifts in phytopathogen- and disease-suppressive soil taxa in agricultural soil microbial communities.^55^ The authors measured paracetamol-induced metabolic activity and proposed these shifts were partially driven by some taxa using paracetamol as a nutrient source.^55^ AS enzymes are known catalyze the first biotransformation step of paracetamol,^56^ thus inviting future research on the role of AS enzymes in wastewater treatment and agricultural dynamics. Within the agricultural context, several known AS enzymes also transform the herbicides, chlorpropham^57,58^ and propanil,^44,58^ In our study, propanil was transformed in our study by eight different AS enzymes. Propanil itself is primarily microbially transformed into 3,4-dichloroaniline, a more toxic and persistent metabolite than the parent compound.^59^

Propanil, paracetamol and other micropollutants included in this study also are known to induce the production reactive oxygen species.^60,61^ This stress response from micropollutants promotes the transfer of antibiotic resistance genes,^62^ giving rise to public health concerns.^63^ A second mechanism promoting antibiotic resistance is the microbial biotransformation of antibiotic conjugates, such as acetylsulfamethoxazole (AcSMX).^64^ Both sulfamethoxazole (SMX) and AcSMX are readily biotransformed in anaerobic urine storage tanks^10^ and wastewater treatment plants.^64^ Four AS enzymes in our study hydrolyzed AcSMX to SMX, reactivating the antibiotic which could lead to inhibitory effects or promotion of antibiotic resistance in wastewater or aquatic microbial communities.^65^

Collectively, these findings suggest AS enzymes may contribute to relevant processes such as microbial community dynamics, contaminant persistence, drug reactivation, and antibiotic resistance. Their global distribution and conservation highlights their relevance in micropollutant biotransformations for agricultural, human, and environmental health.

## Conclusion

In summary, we biochemically characterized a novel AS arylamidase from a *Lacticaseibacillus rhamnosus* urinary isolate and expanded our analysis to 16 additional urine-derived microbial AS enzymes. We identified previously unreported micropollutants as substrates for AS arylamidases, namely pharmaceuticals flutamide, vorinostat, capecitabine, and the antibiotic metabolite acetylsulfamethoxazole. Using a gradient boosting machine learning model, we identified molecular properties as well as surface residues in proximity to the substrate tunnel features predictive of enzyme-substrate specificity. Future studies on these features are warranted to rationally engineer AS enzymes for biocatalysis or bioremediation strategies. Overall, these findings contribute to the development of pharmaceuticals and pesticides that are readily biotransformed, facilitating a society built on ‘benign-by-design’ compounds.

## Data availability

Scripts and data associated with this analysis are available at: https://github.com/MSM-group/amidase-paper

## Acknowledgements

We thank the entire Joint Genome Institute team including Miranda Harmon-Smith, Yasuo Yoshikuni, Ritesh Mewalal, and Ian Blaby. Victoria Poltorak is acknowledged for insightful discussions on machine learning. We thank René Gall for his technical support and Isabelle Teufer and Sophie Kuhn for their experimental assistance. We acknowledge Olga Schubert for idea generation and discussions.

