## Supplementary Information for "Machine learning reveals signatures of promiscuous microbial amidases for micropollutant biotransformations"

### Table of Contents


Supplementary Methods 2

Supplementary Figures 3

Supplementary Tables 15

### Supplementary Methods

#### Substrate specificity screening of P205 with micropollutants

Stock solutions of individual micropollutants were prepared in dimethyl sulfoxide (DMSO) at a concentration of 10 mM and stored at −20°C. These micropollutants were then divided into 14 submixtures, mixed and diluted with ethanol to achieve a final working concentration of 200 µM for each micropollutant.

Micropollutant biotransformations catalyzed by the enzyme P205 were measured in a clear-bottomed 96-well plate (Greiner) as described previously with slight modifications. (1) Briefly, individual micropollutant sub-mixtures were first added to an empty 96-well plate. The plate was placed in a fume hood to allow the organic solvent to completely evaporate. After evaporation, 239 µL of 100 mM Tris-HCl buffer (pH = 8) was added to each well. The plate was placed on a rotary shaker at 50 rpm for 15 min to redissolve the micropollutants. Purified P205 in 11 µL aliquots were added to a 96-well plate to achieve a final enzyme concentration of ~190 nM per well and mixed thoroughly to initiate the biotransformation reactions. Samples were collected at 0 h, 6 h, and 24 h. At each sampling event, a 20 µL sample was taken and mixed with 40 µL pure methanol to completely denature the enzyme and quench the reaction, followed by the addition of 140 µL Tris-HCl buffer to the sample mixture. After centrifugation (13,000 × g, 4°C for 15 min), the supernatant was collected and stored at 4°C for subsequent measurement by ultrahigh-performance liquid chromatography coupled to high-resolution tandem mass spectrometry (UHPLC-HRMS/MS). An abiotic buffer control and inactivated enzyme (P205-S146A) control were set up in the same way as the experiment described above. UHPLC-HRMS/MS analysis was conducted as described below. Matrix match calibration standards were used for quantification. The quantification was based on concentrations calculated from peak area integration using TraceFinder 5.1 (Thermo Fisher Scientific).

#### UHPLC-HRMS/MS analysis

The micropollutants were measured by UHPLC-HRMS/MS (Q Exactive, Thermo Fisher Scientific) using a method as previously described.1 Briefly, for the LC separation, 25 µL samples were injected into an ACQUITY Premier BEH C18 Column (particle size 1.7 µm, 100 × 2.1 mm, Waters) and eluted with Nanopure water (A) and methanol (B), both containing 0.1% formic acid, at a flow rate of 300 µL/min. A linear elution gradient was set as follows: 95% A: 0 – 1.5 min, 95% – 5% A:1 – 7.5 min, 5% A: 7.5 – 9.5 min, and 95% A: .5 – 11.5 min. For HRMS, mass spectra were acquired in full scan mode at a resolution of 70,000 at m/z 200 and a scan range of m/z 100 − 1000 in positive/negative switching electrospray ionization mode. Xcalibur 4.0 (Thermo Fisher Scientific) and TraceFinder 5.1 (Thermo Fisher Scientific) were used for data acquisition and analysis. Compounds exhibiting more than 40% parent compound removal by P205 and negligible removal in abiotic and inactivated (P205-S146A) controls were selected for further investigation.

### Supplementary Figures

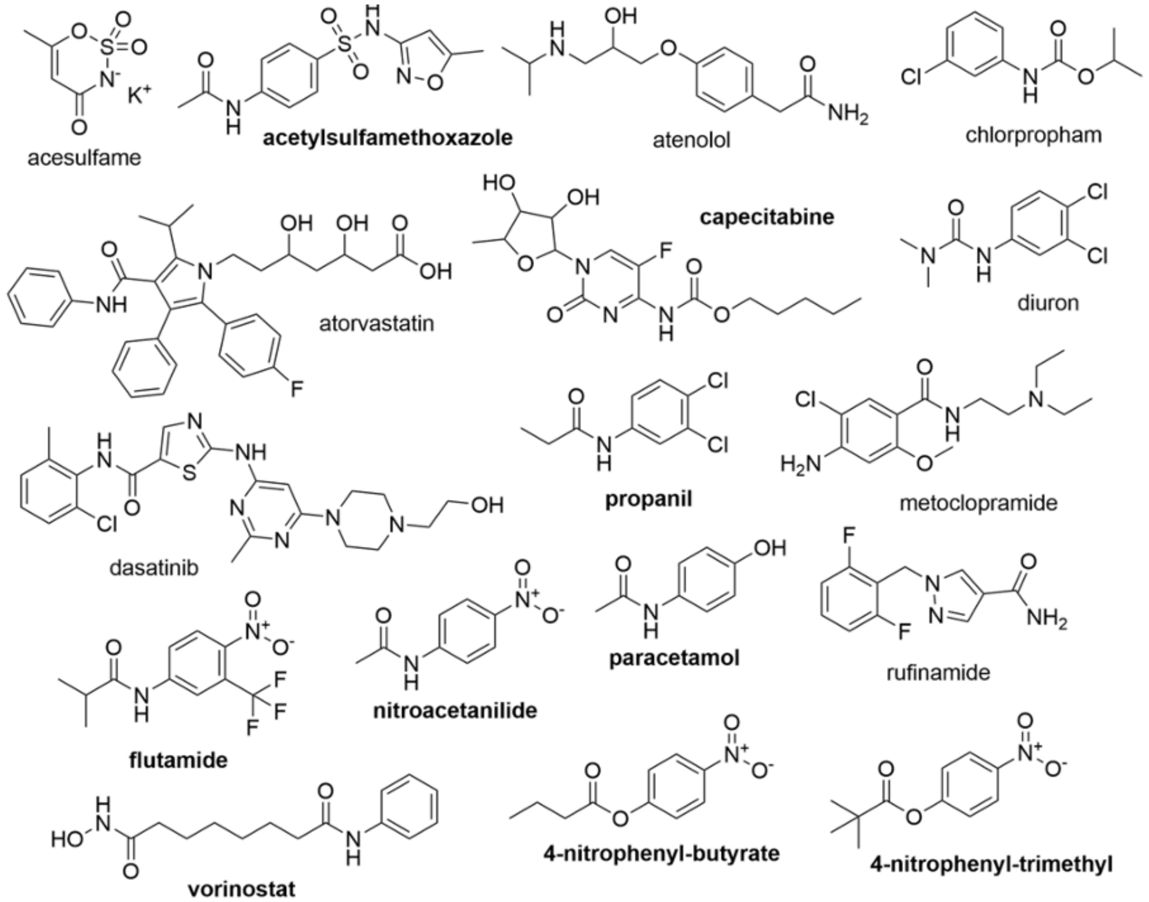


**Supplementary Figure 1**: Chemical structures of the substrates tested in this study against the AS enzyme library. Bold names indicated substrates that were biotransformed by at least one enzyme.

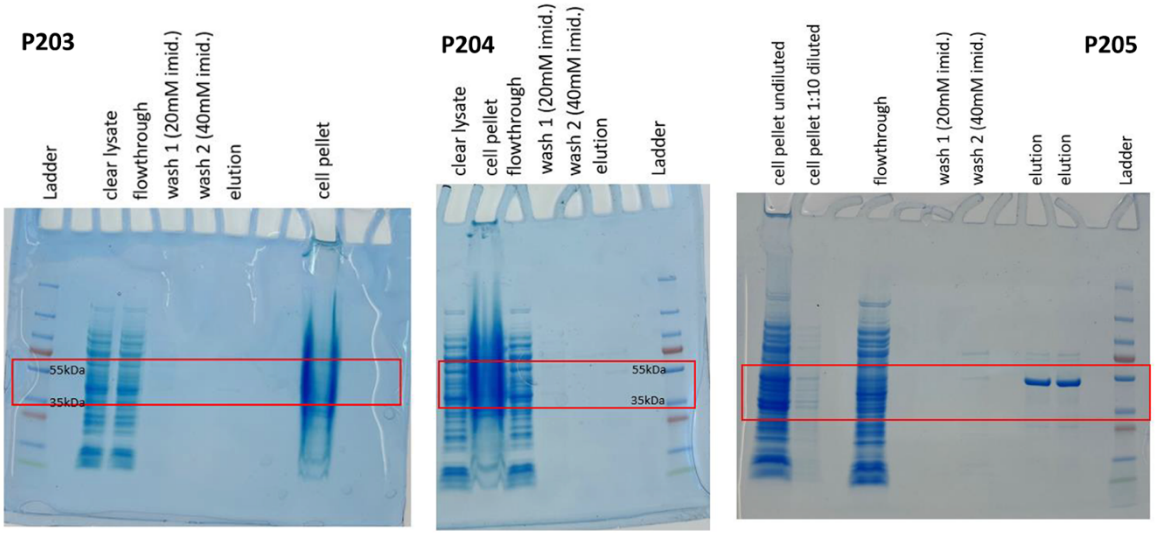


**Supplementary Figure 2:** SDS-PAGE gels of protein purification of P203, P204 and P205 via his-tag affinity chromatography. Red boxes show protein bands between sizes of 35 to 55 kilodaltons (kDa).

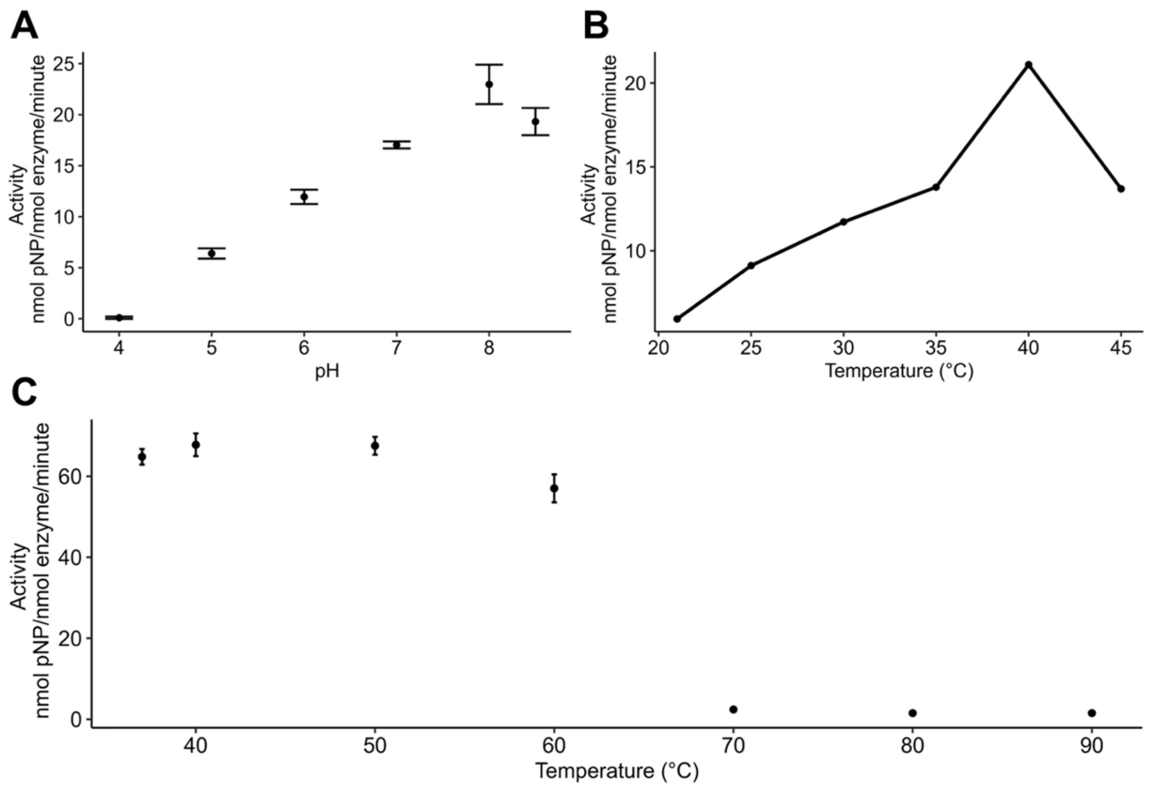


**Supplementary Figure 3:** Biochemical characterization of P205. **A)** The P205 pH optimum was determined to be pH 8.0 by measuring the isosbestic point of 4-nitrophenol (347 nm) across pH 4-9 with the substrate 4-nitrophenyl trimethylacetate. Citrate buffer was used for pH 4-6 and Tris-HCl buffer was used for pH 7-9. Enzyme activity measurements for pHs higher than 9 are not shown due to rapid abiotic hydrolysis of 4-nitrophenyl trimethylacetate. **B)** The P205 temperature optimum was determined measuring 4-nitrophenol absorbance at 410 nm in Tris-HCl buffer (pH 8) across different temperatures using 4-nitrophenyl-acetate. **C)** The heat stability of P205 was assessed by measuring 4-nitrophenol absorbance at 410 nm in Tris-HCl buffer (pH 8) after exposure to various temperatures for 1 hour using 4-nitrophenyl-butyrate. All experiments were conducted using purified P205 (final concentration of 0.0022 mg/mL for the temperature optimum, 0.022 mg/mL for the pH optimum and heat stability).

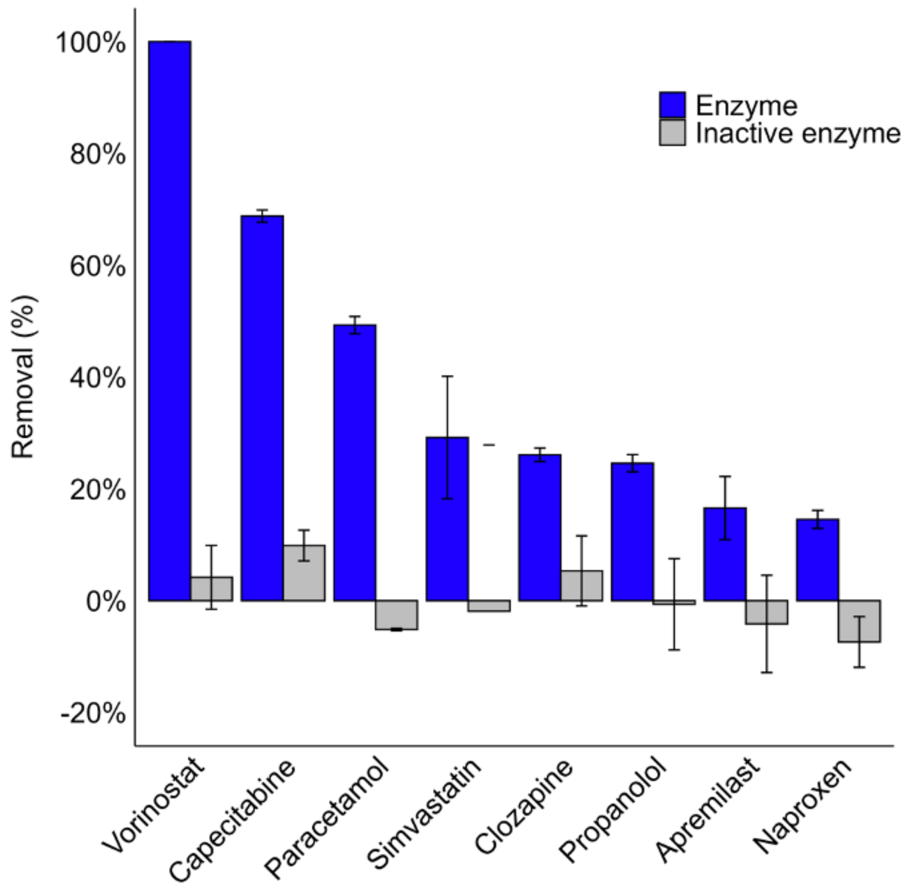


**Supplementary Figure 4:** P205 (blue) substrate specificity screening hits (with >20% removal difference between enzyme and inactive enzyme samples) after 24 h incubation with a collection of a library of 183 micropollutants. The inactive enzyme control (gray) is the catalytically inactive variant P205-S146A.

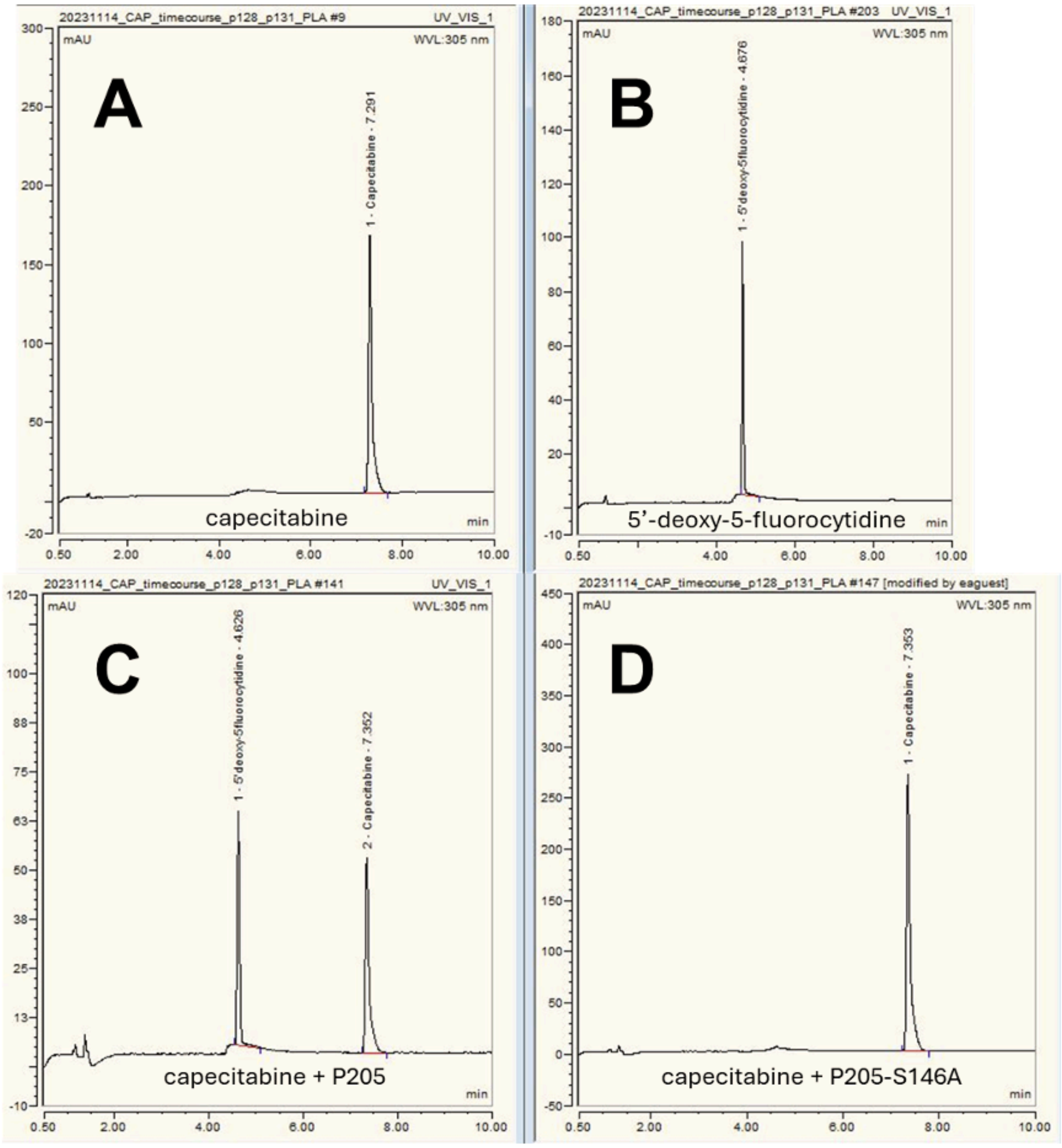


**Supplementary Figure 5**: HPLC chromatograms and 3D spectra of **A)** capecitabine (RT 7.4 min) **B)** an authentic standard of the transformation product 5'-deoxy-5-fluorocytidine (RT 4.6 min), and **C)** capecitabine incubated with purified P205 (0.05 mg/mL) for 8 hours, and **D)** capecitabine incubated with purified P205-S146A (0.05 mg/mL) for 8 hours.

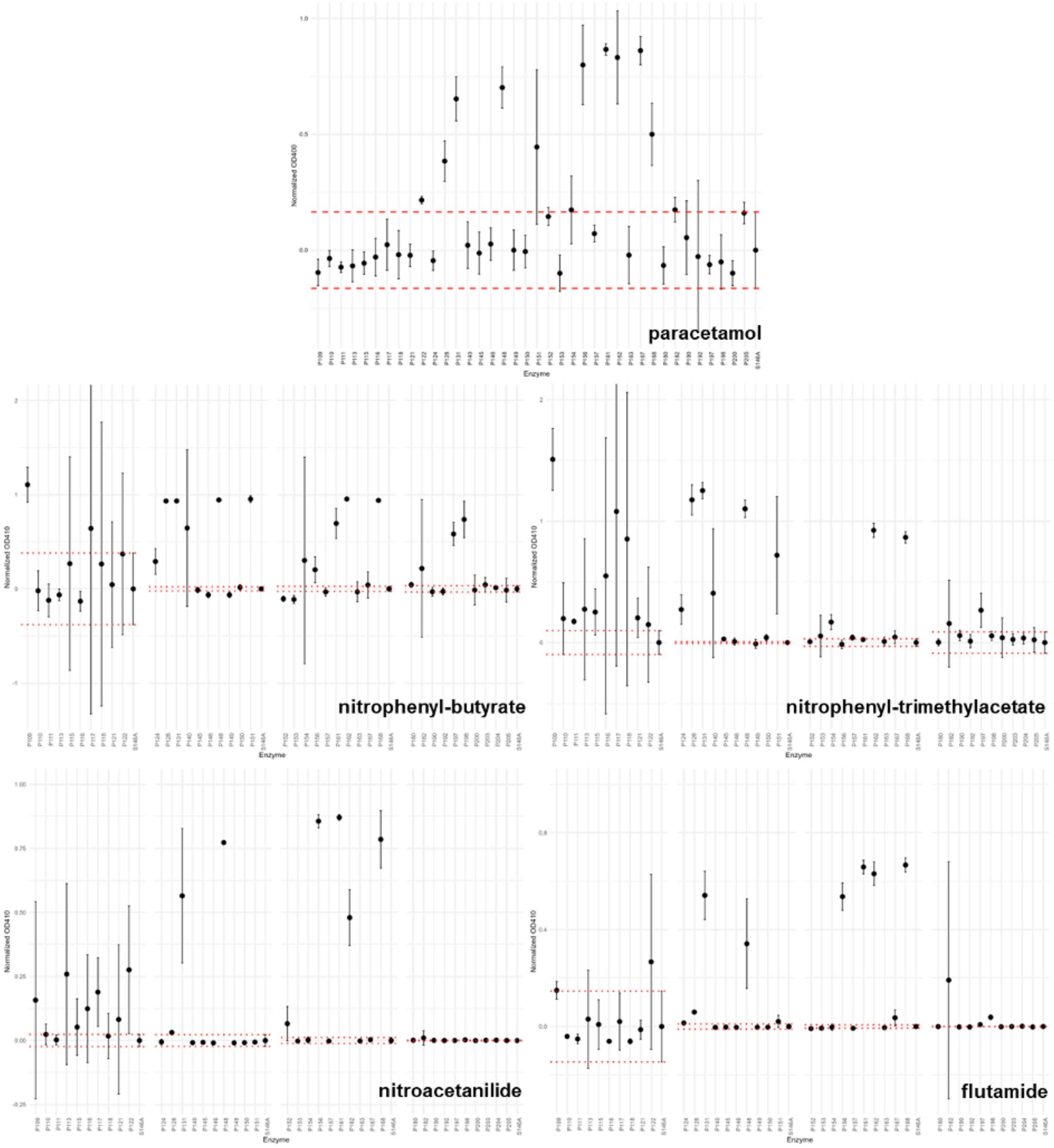


**Supplementary Figure 6:** Endpoint ODs measuring the absorbance of chromophores produced by the activity of the AS enzyme library (see Methods). The OD was normalized by subtracting the OD of the inactivated enzyme control (P205-S146A, depicted as S146A). The red dotted lines represent two standard deviations from the mean activity of P205-S146A. Datapoints are from biological triplicates.

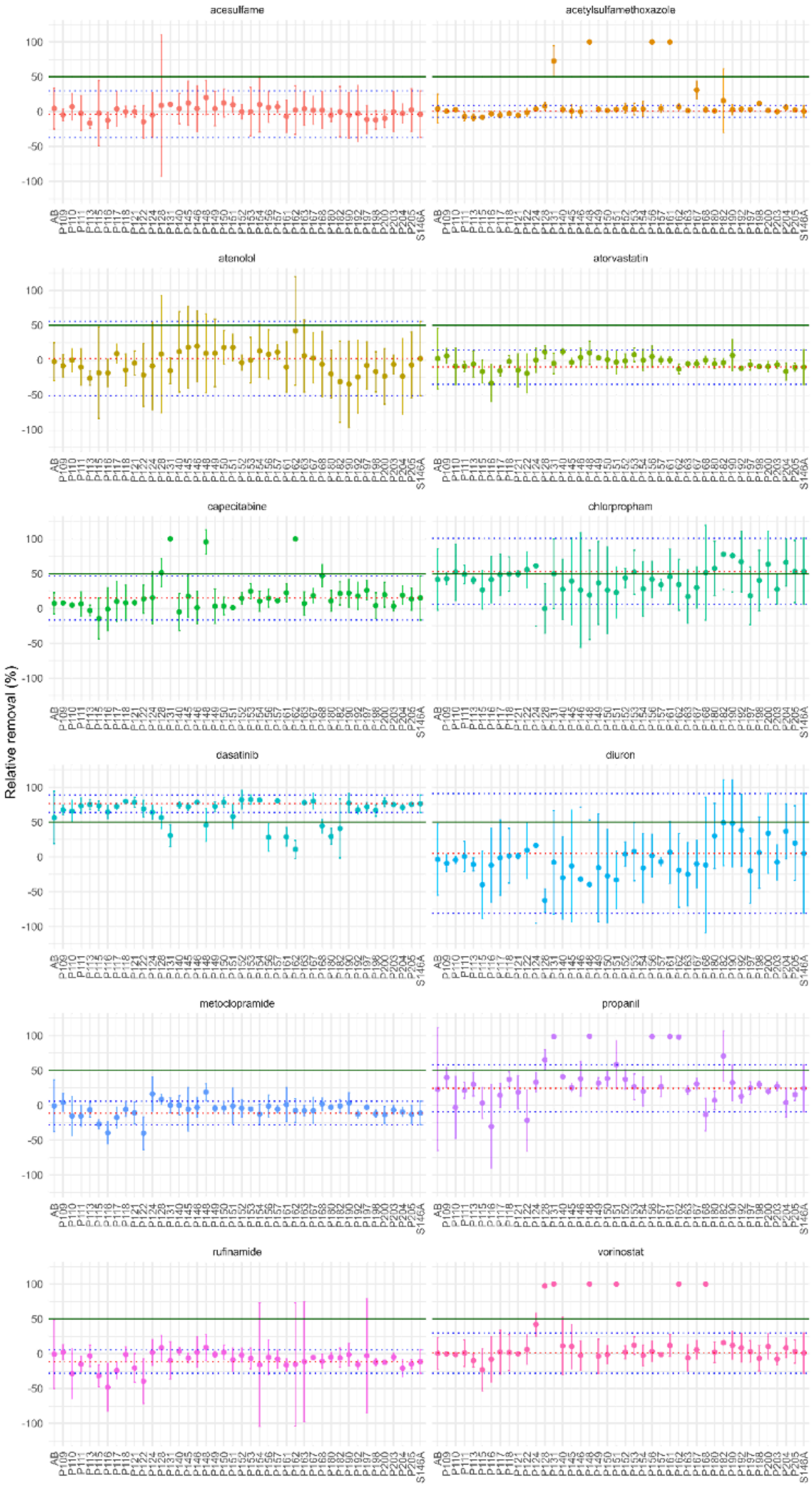
acesulfame
acetylsulfamethoxazole
atenolol
atorvastatin
capecitabine
chlorpropham
dasatinib
diuron
metoclopramide
propanil
rufinamide
vorinostat
Relative removal (%)
100
50
0
-50
-100

**Supplementary Figure 7**: Relative removal of substrates by AS enzyme library. The green line marks the 50% removal threshold (first criterion for activity). AB represents the abiotic control, and S146A denotes the catalytically inactive variant P205-S146A. The dotted red line indicates the median of P205-S146A, while the blue lines represent the 1.5 IQR values of P205-S146A. Values above the media + 1.5 IQR of P205-S146a were considered to meet the second criterion for activity. These stringent criteria were selected to filter out false positives due to abiotic removal (e.g., chlorpropham, dasatinib). Different wavelengths were used for quantification: 225 nm (acesulfame, atenolol, chlorpropham, rufinamide), 252 nm (propanil, diuron, atorvastatin, acetylsulfamethoxazole, vorinostat), and 305 nm (dasatinib, metoclopramide, capecitabine). Datapoints are from biological triplicates.

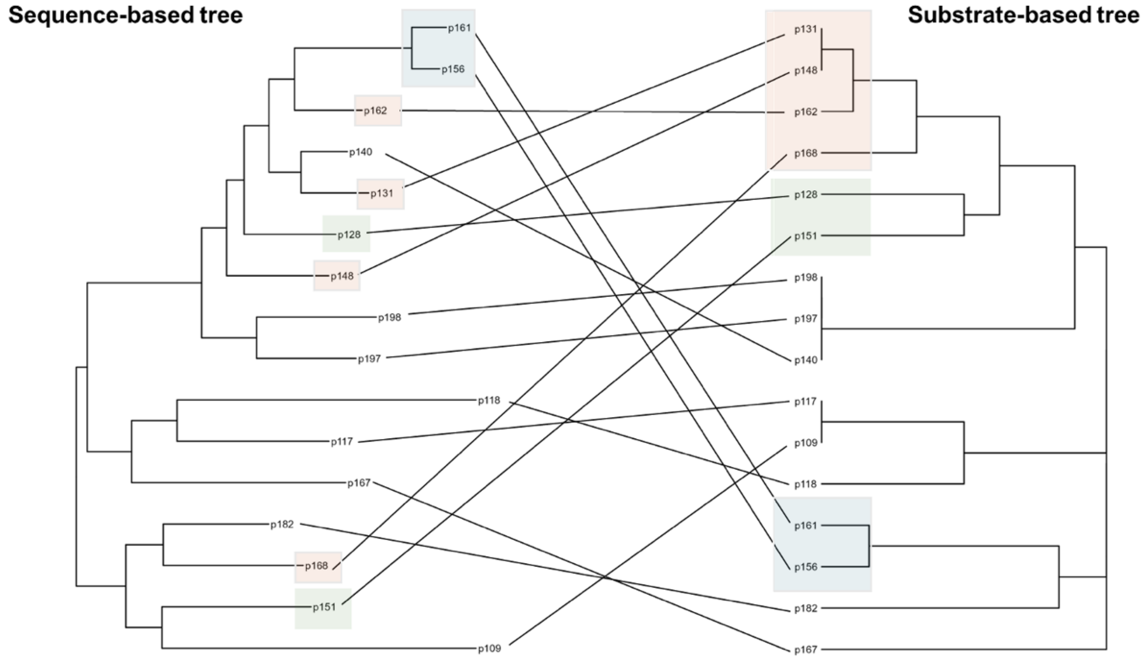


**Supplementary Figure 8**: Substrate specificity dendrogram (excluding substrates that are not biotransformed by any enzyme) based on Jaccard index calculated from the substrate preference matrix. Colored boxes indicate enzymes active with at least 4 substrates.

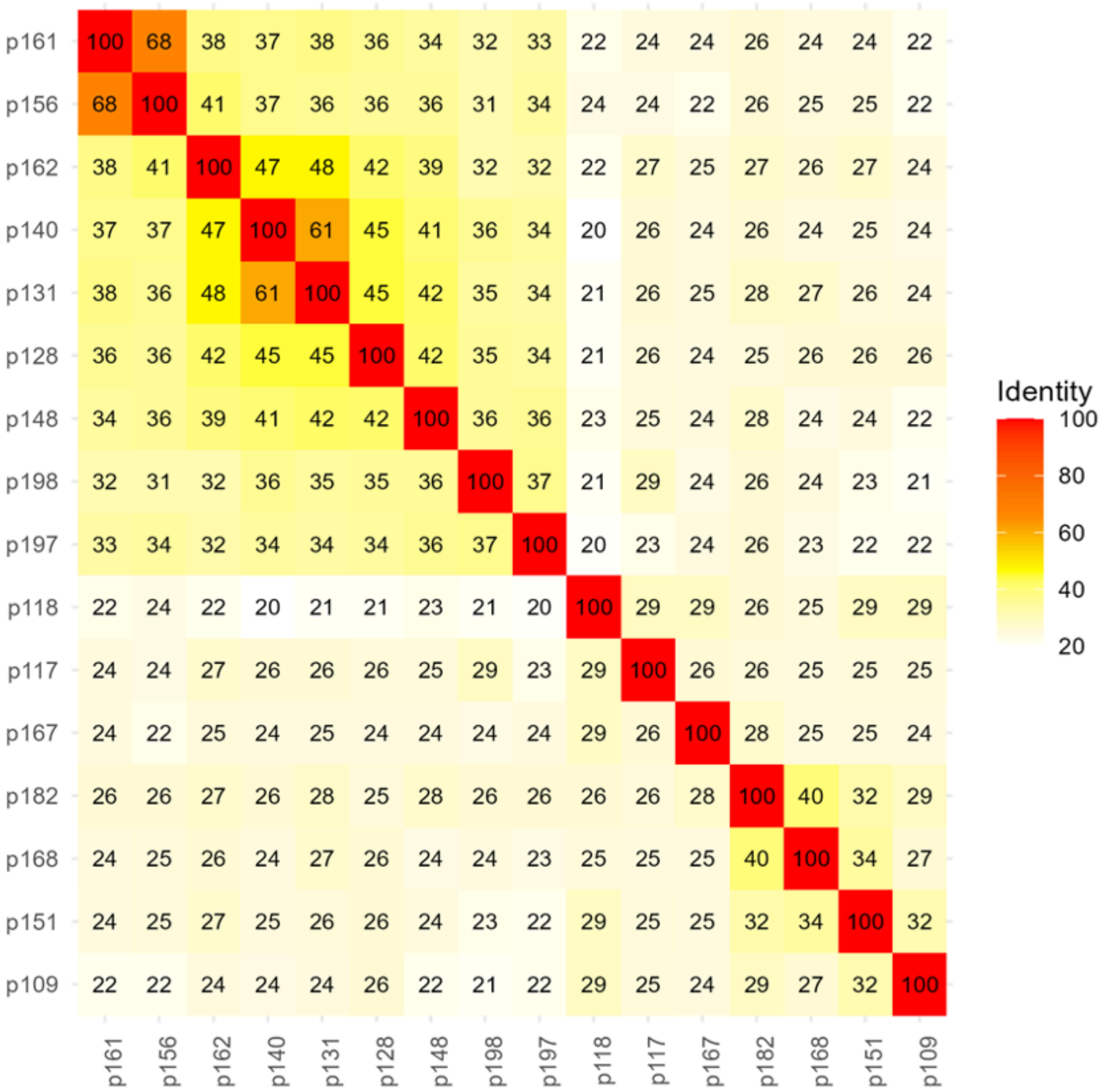


**Supplementary Figure 9**: Protein identity matrix of the 16 enzymes which showed activity with at least one substrate.

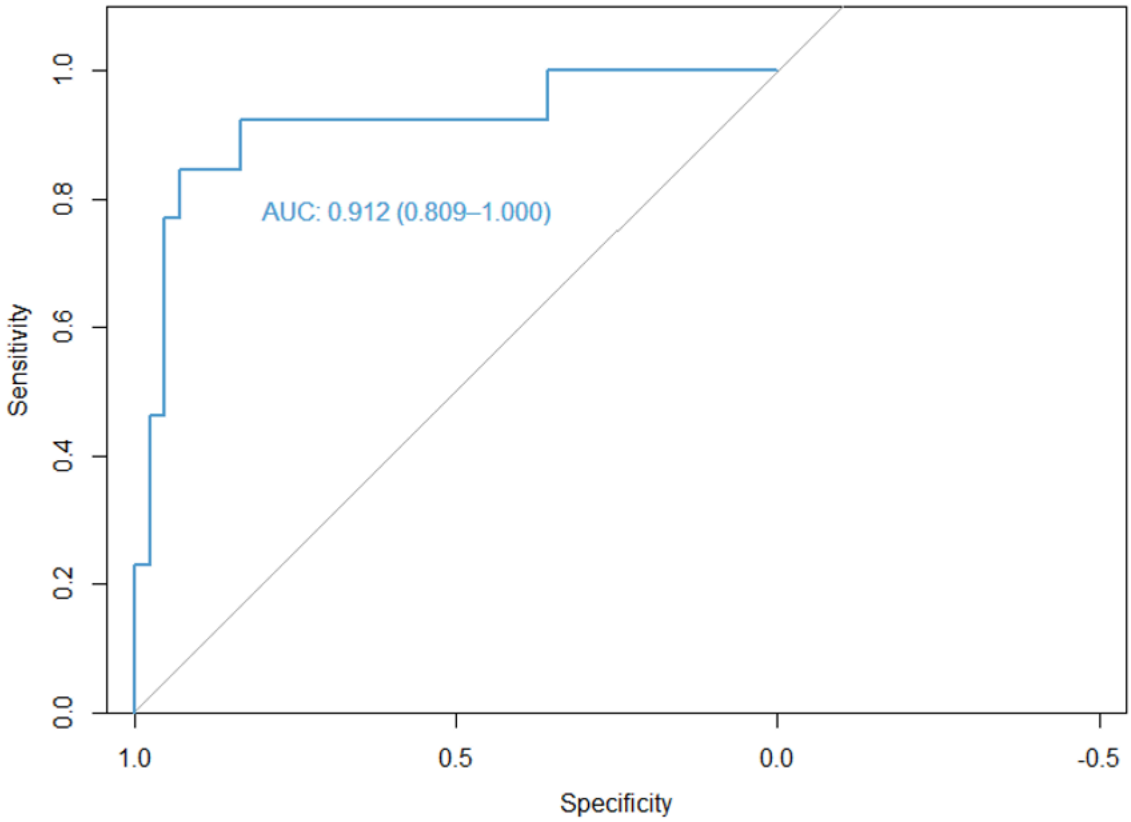


**Supplementary Figure 10**: Receiver operating characteristic (ROC) curve of the gradient boosting model trained on the 272 enzyme-substrate pairs using featurization of chemical functional groups and amino acids positions of the amidase signature region.

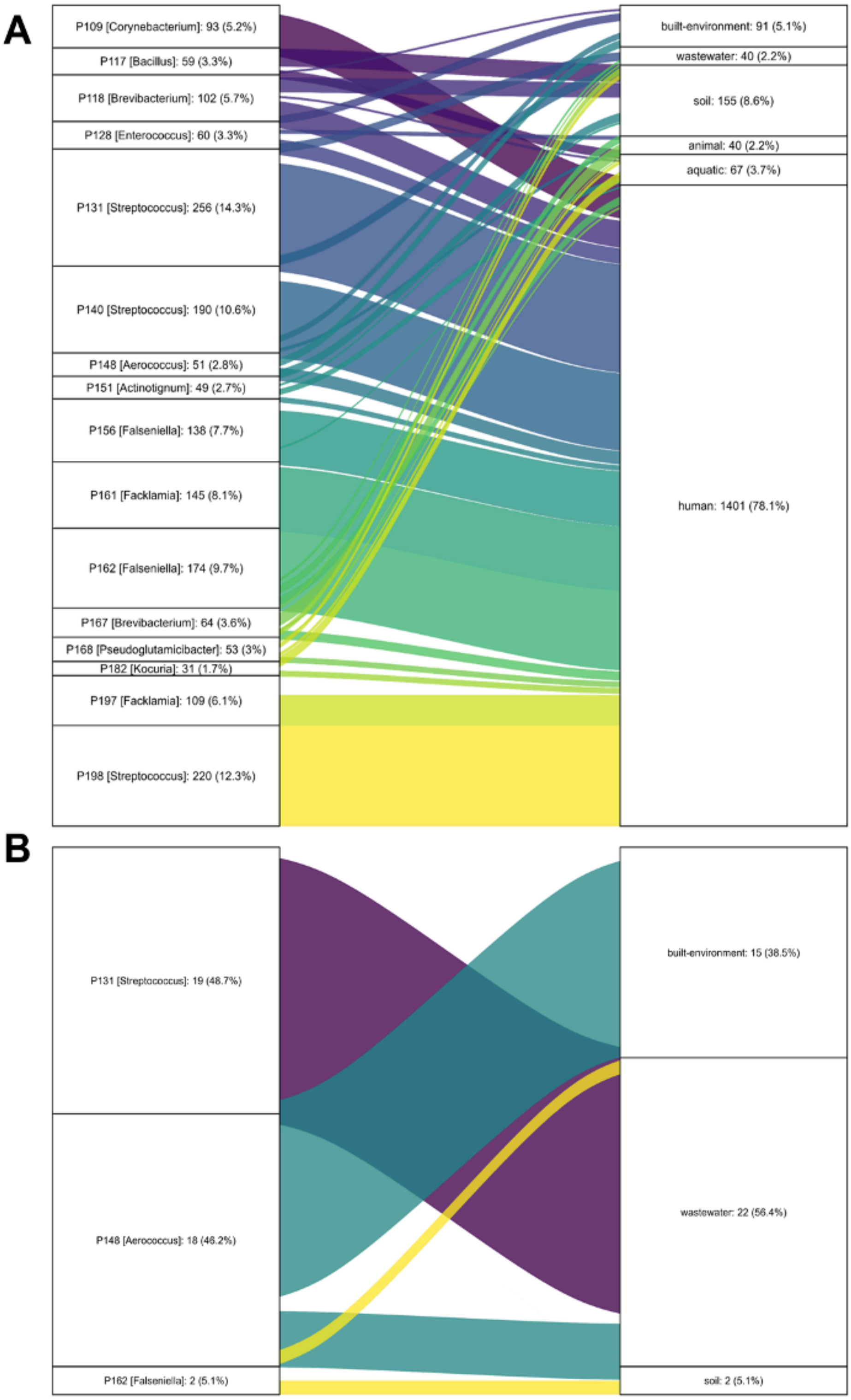


**Supplementary Figure 11:** Distribution of enzyme homologs across different environments (according to the Global Microbial Gene Catalog). **A**) Prevalence of active enzyme homologs in environmental, animal and human microbiomes. **B**) Prevalence of the most promiscuous enzymes in the study in environmental microbiomes. Each stratum represents an enzyme, with the bacterial host genus indicated in brackets. The connections between strata represent the prevalence of each enzyme in various environments. The labels on the strata display both the absolute counts and the relative proportions of the different enzyme homologs.

### Supplementary Tables

**Supplementary Table 1:** Substrates and compounds used in this study

| Compound | Supplier |
|---|---|
| acesulfame | Sigma-Aldrich |
| atenolol | Chemie Brunschwig |
| acetylsulfamethoxazole | Chemie Brunschwig |
| atorvastatin | Chemie Brunschwig |
| capecitabine | Chemie Brunschwig |
| chlorpropham | Chemie Brunschwig |
| diuron | Chemie Brunschwig |
| dasatinib | Sigma Aldrich |
| metoclopramide | Chemie Brunschwig |
| propanil | Sigma Aldrich |
| rufinamide | Chemie Brunschwing |
| vorinostat | Chemie Brunschwig |
| 4-nitrophenyl butyrate | Sigma-Aldrich |
| 4-nitrophenyl trimethylacetate | Sigma-Aldrich |
| flutamide | Sigma-Aldrich |
| nitroacetanilide | Sigma-Aldrich |
| paracetamol | Sigma-Aldrich |
| 4-nitrophenol | Sigma-Aldrich |
| nitroaniline | Sigma-Aldrich |
| 4-nitro-3-(trifluoromethyl)aniline | Sigma Aldrich |

**Supplementary Table 2:** Solutions for protein purification using benchtop purification

| | |
|---|---|
| Equilibration buffer | 20 mM Tris-HCl, 500 mM NaCl, 10% glycerol, 20 mM imidazole, pH 8 |
| Wash buffer | 20 mM Tris-HCl, 500 mM NaCl, 10% Glycerol, 40 mM imidazole, pH 8 |
| Elution buffer | 20 mM Tris-HCl, 500 mM NaCl, 10% Glycerol, 500 mM imidazole, pH 8 |
| SGT buffer | 5 mM Tris-HCl, 30 mM NaCl, 10 % glycerol, pH = 8.0 |

**Supplementary Table 3:** Shorthand notation, NCBI accession numbers, and source organisms for the amidase signature enzymes investigated in this study.

| **Shorthand notation** | **Accession number** | **Source organism** |
|---|---|---|
| P109 | PLA27924.1 | *Corynebacterium coyleae* |
| P110 | PKZ65245.1 | *Gordonia terrae* |
| P111 | PLA20978.1 | *Corynebacterium pyruviciproducens* |
| P113 | PKZ43269.1 | *Globicatella sanguinis* |
| P115 | PMC16657.1 | *Oligella urethralis* |
| P116 | PLA35052.1 | *Corynebacterium amycolatum* |
| P117 | PLR73545.1 | *Bacillus* sp. |
| P118 | PKY70760.1 | *Brevibacterium ravenspurgense* |
| P121 | PLB08528.1 | *Pseudomonas aeruginosa* |
| P122 | PLA16723.1 | *Klebsiella pneumoniae* |
| P124 | PLA23573.1 | *Cutibacterium acnes* |
| P128 | PKZ01497.1 | *Enterococcus faecalis* |
| P131 | PKZ94153.1 | *Streptococcus salivarius* |
| P140 | PLA78093.1 | *Streptococcus agalactiae* |
| P145 | PKY92026.1 | *Aerococcus christensenii* |
| P146 | PLR70916.1 | *Bacillus* sp. |
| P148 | PKZ20744.1 | *Aerococcus sanguinicola* |
| P149 | PKZ65816.1 | *Gordonia terrae* |
| P150 | PLT14322.1 | *Limosilactobacillus fermentum* |
| P151 | PLB82905.1 | *Actinotignum timonense* |
| P152 | PLB04707.1 | *Pseudomonas aeruginosa* |
| P153 | PLR66227.1 | *Bacillus* sp. |
| P154 | PMC29638.1 | *Lactobacillus iners* |
| P156 | PKY89575.1 | *Falseniella ignava* |
| P157 | PLA40941.1 | *Neisseria sicca* |
| P161 | PKY92403.1 | *Facklamia hominis* |
| P162 | PKY89520.1 | *Falseniella ignava* |
| P163 | PKZ08631.1 | *Dermabacter hominis* |
| P167 | PKY70105.1 | *Brevibacterium ravenspurgense* |
| P168 | PKY79758.1 | *Pseudoglutamicibacter albus* |
| P180 | PKZ05429.1 | *Enterococcus faecalis* |
| P182 | PKZ37694.1 | *Kocuria rhizophila* |
| P190 | PKZ63448.1 | *Gordonia terrae* |
| P192 | PKY80781.1 | *Pseudoglutamicibacter albus* |
| P197 | PKY93381.1 | *Facklamia hominis* |
| P198 | PLA79535.1 | *Streptococcus agalactiae* |
| P200 | PKY92027.1 | *Aerococcus christensenii* |
| P203 | PKZ25206.1 | *Corynebacterium aurimucosum* |
| P204 | PKZ66103.1 | *Gordonia terrae* |
| P205 | PLA55254.1 | *Lacticaseibacillus rhamnosus* |
